# Integration of short- and long-term responses to environmental stimuli shape seasonal transcriptome dynamics

**DOI:** 10.1101/2021.08.02.454700

**Authors:** Yuko Kurita, Hironori Takimoto, Mari Kamitani, Yoichi Hashida, Makoto Kashima, Ayumi Tezuka, Takanari Tanabata, Atsushi J. Nagano

## Abstract

Plants must respond to various seasonally changing environmental stimuli. In a previous study, seasonally oscillating genes were identified by a massive time-series transcriptome analysis for the wild population of *Arabidopsis halleri* ssp. *gemmifera*, a sister species to *Arabidopsis thaliana*. It was not clear how environmental stimuli shaped the seasonal expression pattern of these seasonally oscillating genes. In this study we show that responses in different time-scales contributed to the formation of seasonal expression patterns for several genes. To analyze the seasonally oscillating genes, we established an experimental system to mimic seasonal expression trends using *A. thaliana* and a “smart growth chamber mini,” a hand-made low-cost small chamber. *Arabidopsis thaliana* plants were cultured under conditions that mimicked the average monthly temperatures and daylengths under different day-scale incubation. In total, the seasonal trends of 1627 seasonally oscillating genes were mimicked, and they showed varying temporal responses (constant, transient, and incremental) to environmental stimuli. Our results suggest that plants perceive and integrate information regarding environmental stimuli in the field by combining seasonally oscillating genes with different temporal responsiveness.

## Introduction

Plants responses to seasonal changes in environmental factors are essential mechanisms. The analysis of gene expression patterns is a useful approach to understand how plants respond to environmental stimuli and adapt physiologically and morphologically. Over the past few decades, a number of studies have analyzed seasonal gene expression patterns (Andersson et al., 2004; Galindo González et al., 2012; Lu et al., 2020; Nagano et al., 2019, 2012). Vigorous research has uncovered the mechanisms of several seasonal phenomena, such as vernalization in flowering, growth cessation, and bud dormancy (Falavigna et al., 2019; Luo and He, 2020; Maurya and Bhalerao, 2017; Song et al., 2013). However, there are many seasonal phenomena whose mechanisms have not yet been elucidated, or genes that show seasonally oscillating expression patterns but whose functions have not yet been clarified.

The previous study have identified several genes that show seasonal oscillation patterns in gene expression in *Arabidopsis halleri* ssp. *gemmifera* (*Ahg*), a perennial herbaceous plant closely related to *Arabidopsis thaliana* (*Ath*) by seasonal transcriptome analysis (Nagano et al., 2019). The seasonal expression patterns of many of these seasonally oscillating genes (SO genes) were defined by annual temperature changes rather than annual changes in daylength. However, the molecular mechanisms regulating the seasonal expression of a large number of SO genes are still unknown. Expression patterns of several SO genes are thought to be affected by long-term environmental stimuli (month-scale response: weeks to months) such as *FLC*, which is associated with vernalization in *Ath* (Bouché et al., 2017; Jean Finnegan, 2015). On the contrary, it is thought that other SO genes show a minute-scale response (minutes to hours) to the environmental stimuli, such as *CBF/DREB1s* for the cold stress response (Gong et al., 2020; Ye et al., 2019). In addition, the mechanism of day-scale response (1 day to 1 week) to repeated temperature stimulation has recently attracted attention (Leuendorf et al., 2020; Yamaguchi et al., 2021). Nagano et al. revealed that there are numerous genes whose expression can be explained by meteorological data from the past one to several days (Nagano et al., 2012).

The seasonal phenomena of plants are fascinating; however, field studies are time-consuming and challenging. Moreover, in the field, plants are simultaneously exposed to many biotic and abiotic environmental stimuli. To understand the molecular mechanisms of plant responses in the field, effective experimental systems which can separate and control the environmental stimuli, as well as field studies, are essential. Recently, combination studies of the field and experimental systems, such as ecotrons (Roy et al., 2021) and the smart growth chamber (SGC) (Hashida et al., 2022), which can reproduce microclimatic conditions, have been reported. In the study of the function of SO genes, understanding their properties by using controlled environments will be helpful. For example, what stimuli shape the seasonal trends of the SO gene and the length of time required for stimulus acceptance.

In this study, we focused on the responses to temperature and day-length occurring within several days (day-scale responses). We cultured *Ath* plants under simplified monthly conditions with average temperature and daylength of each month for 7, 3, or 1-day(s) to test how many genes can mimic the seasonal expression of *Ahg*. Compared to *Ahg* plants, which are difficult to culture in large quantities in the laboratory, *Ath* plants can be cultured easily and quickly, and abundant genomic information of such plants is available. In addition, *Ath* allows for further research on SO genes using a variety of molecular genetic tools. To culture plants in parallel under different temperature and daylength conditions, we developed a hand-made low-cost small chamber, called a smart growth chamber mini (SGCmini).

## Results

### Development of a space- and cost-saving incubator with parallel control

To cultivate plants under various conditions, we developed a space- and cost-saving small incubator named as “smart growth chamber mini” (SGCmini) (Fig. **1A**; Supplementary Fig. **S1**). The SGCmini can control the inside temperature within a range of −9 to +20 °C from the outside temperature. The Peltier unit was controlled to equalize the average of the measured temperatures of the two sensors installed at the diagonal corners of the incubator body with the set temperature (Fig. **1B–G**). It took a few minutes to 30 min for the average temperature in the SGCmini to reach the set temperature (Fig. **1C**,**D**). The temperature records of the SGCmini under the March and August conditions are shown in Fig. **1E** and **1F**. Multiple SGCminis can be controlled using a single PC. This allows us to save space and conduct multiple experiments under various conditions simultaneously in one room.

**Fig. 1.**
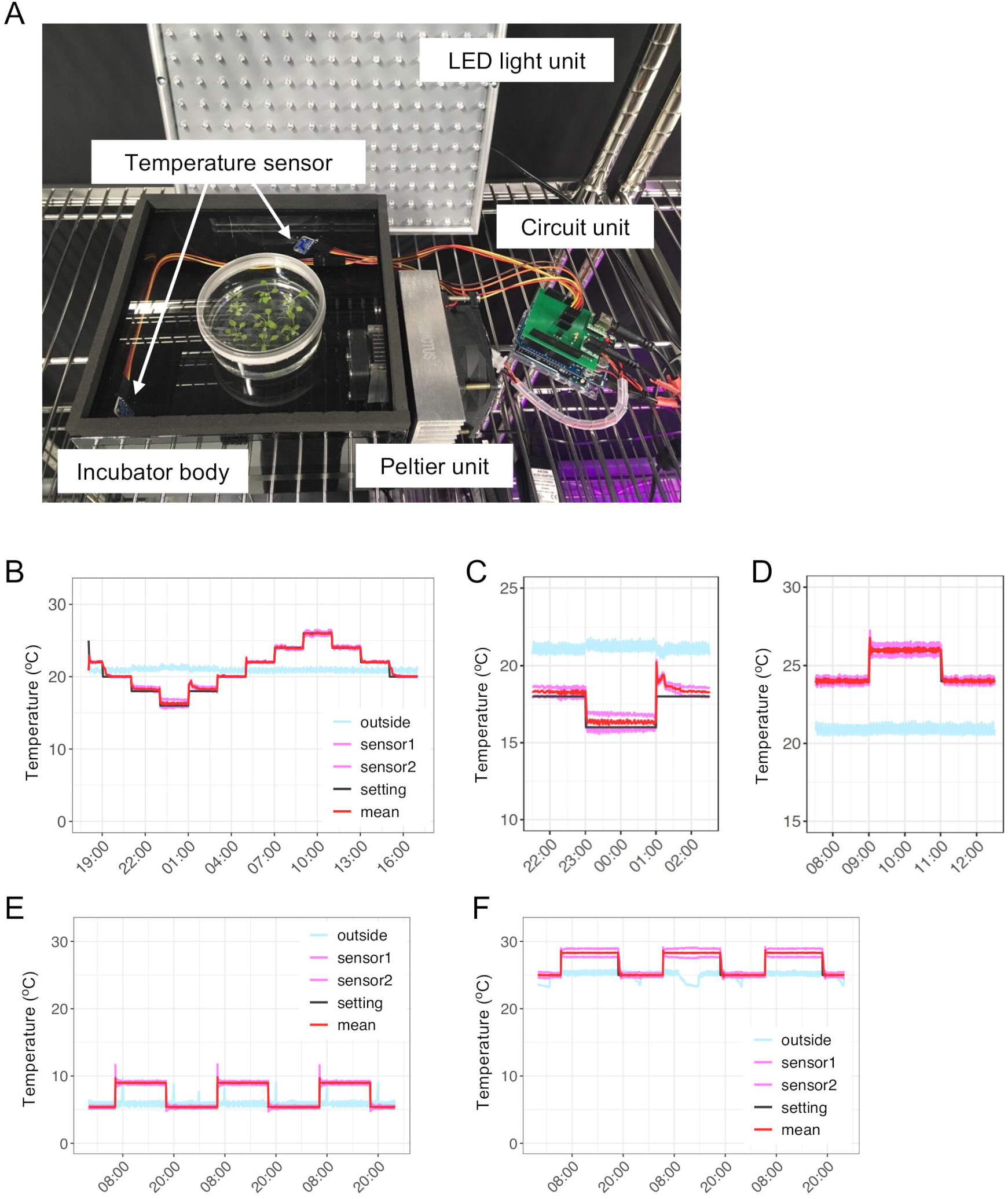
Performance of the smart growth chamber mini (SGCmini). (A) Picture of the SGCmini. The walls of the incubator body were painted for shading. (B) Temperature plot of a test run. The temperature was changed stepwise from 16 °C to 26 °C every 2 h. Light blue line: outside temperature; pink lines: inside temperature measured by the sensors; black line: preset temperature; red line: average temperature of sensor 1 and sensor 2. (C) Enlarged plot of (B) at the lowest temperature setting. (D) Enlarged plot of (B) at the highest temperature setting. (E) Temperature plot for 3 d under the March condition. (F) Temperature plot for 3 d under the August condition.

### The 7-d culture in smart growth chamber mini induced morphological differences under conditions of each month

To mimic the seasonal trend of gene expression in *A. halleri* (*Ahg*) in the natural environment, *A. thaliana* (*Ath*) plants were cultivated in the SGCmini for 7 d under simplified temperature and daylength settings based on the conditions of the natural habitat for each month (Supplementary Table **S2**; Fig. **2A,B**). In the Northern Hemisphere, summer occurs from June through August, and winter occurs from December through February. Sunrise and sunset times on the 15^th^ of each month were set as light-on and light-off times. The average temperature from sunrise to sunset for each month was set as the daytime temperature, and the average temperature from sunset to sunrise for each month was set as the nighttime temperature.

**Fig. 2.**
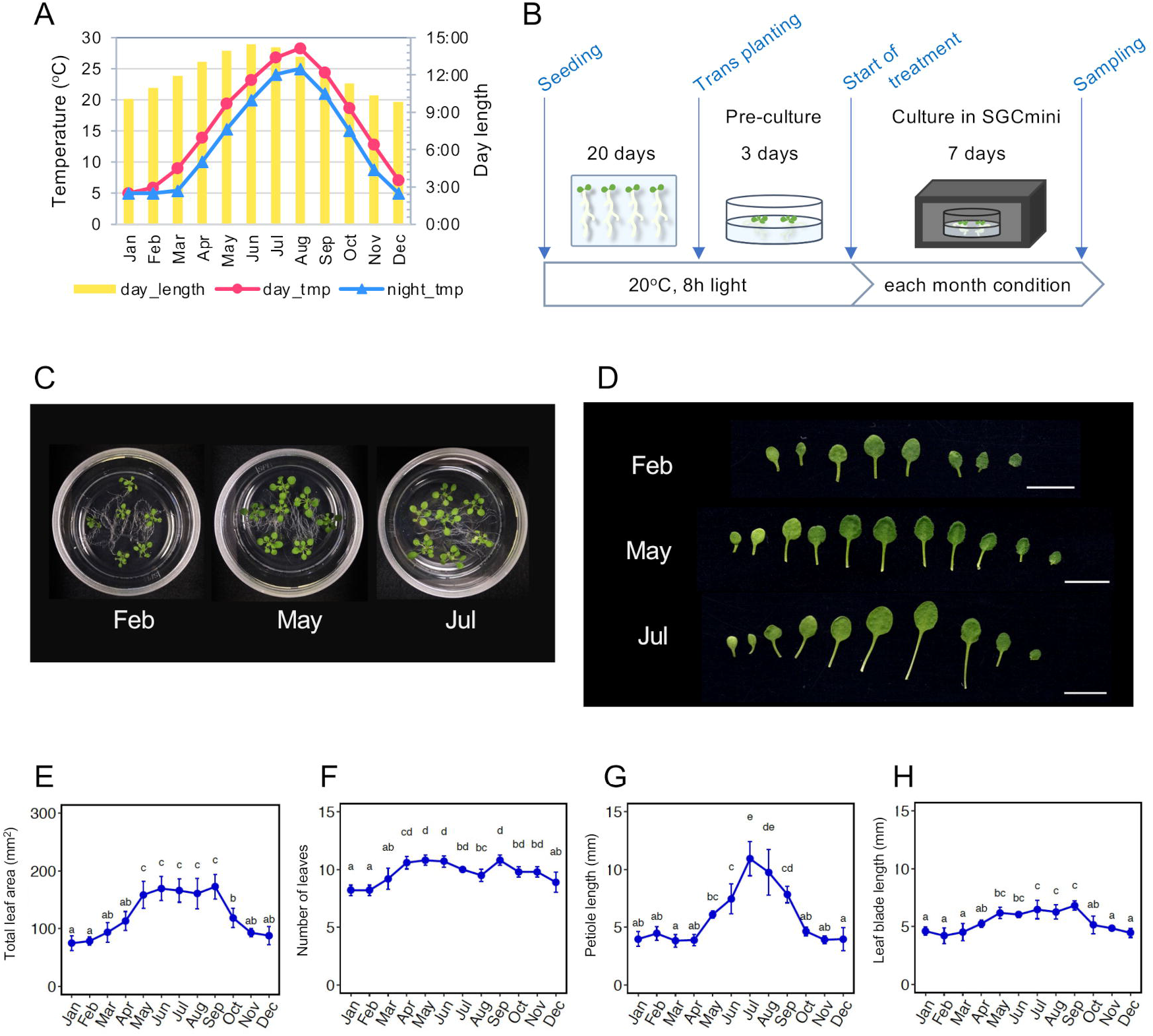
Experimental settings and morphological traits of *Arabidopsis thaliana* (*Ath*) plants grown in the SGCmini for 7 d. (A) Setting of each condition in the SGCmini. Yellow bars: daylength; red circles: daytime temperatures; blue triangles; nighttime temperatures. (B) Schematic diagram of the culture condition. (C) Whole images of plants grown under February, May, and July conditions. (D) Leaves of plants grown under February, May, and July conditions. The youngest leaves are on the right. Bar = 10 mm. (E) Total leaf area. (F) Number of leaves. (G) Maximum petiole length. (H) Maximum leaf blade length. Bar: SD. n = 5–9. Different letters indicate differences between conditions at *P* < 0.05 using ANOVA and Tukey’s HSD test.

The total leaf area and maximum leaf blade length increased from February to May, remained constant from May to September, and decreased after October (Fig. **2C–E,H**). The number of leaves increased from February to May, remained constant from May to June, decreased slightly in July and August, and then increased again in September before decreasing after October (Fig. **2F**). The petiole length increased from April to July and then decreased after August (Fig. **2D,G**). Thus, the 7-d culture induced morphological differences under each month’s condition.

### Seasonal expression of *Ahg* was partially reproduced in *Ath* plants cultured for 7 days under the condition of each month

To analyze how many genes mimic the seasonal expression trend of *Ahg* genes, gene expression data of *Ath* plants grown in the SGCmini for 7 d were obtained by RNA-Seq. Samples with more than 10^5.5^ reads were used for analysis (*Ahg*: 474 samples; *Ath*: 72 samples; Supplementary Fig. **S2A**). *Ahg*-expressed genes (14587 genes) (log_2_(mean rpm +1) > 2) and their orthologous genes in *Ath* were used for analysis (Supplementary Fig. **S2**). The seasonal trend of gene expression was summarized as the amplitude (*α*) and phase (*φ*) by cosine curve fitting (Fig. **3A**). The phase was defined as the time point of the highest gene expression in a year. We defined genes whose amplitude was greater than 1 as SO gene (Fig. **3B**).

**Fig. 3.**
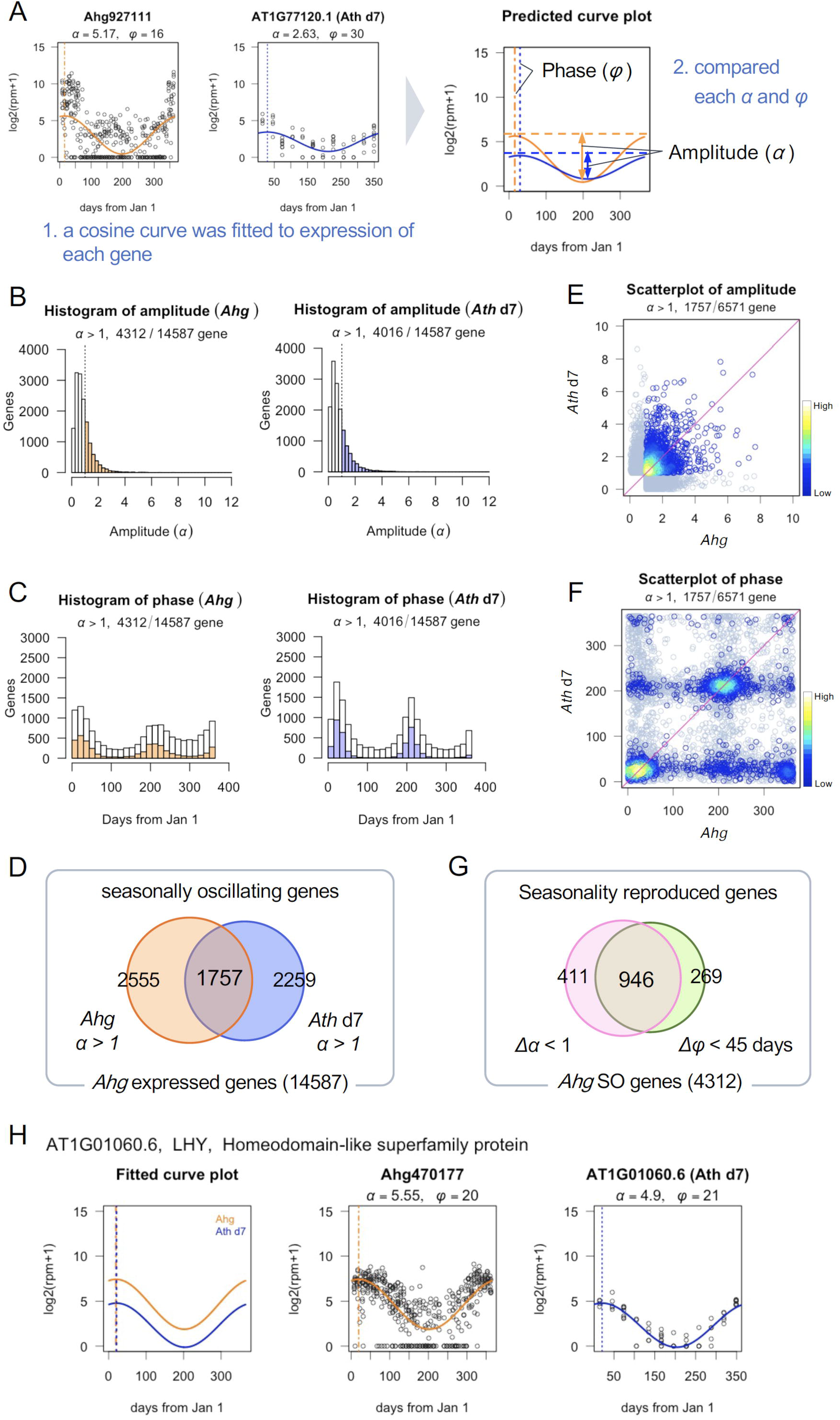
Comparison of seasonal trends of gene expression in *Arabidopsis halleri* ssp. *gemmifera* (*Ahg*) in the field and *Arabidopsis thaliana* (*Ath*) in the SGCmini for 7 d. (A) Schematic diagram of comparison method of seasonal trends of *Ahg* and *Ath*. Black circles indicate observed value by RNA-Seq. The solid orange (*Ahg*) and blue (*Ath*) lines show the fitted cosine curve. The orange and blue dashed lines indicate the phases of *Ahg* and *Ath*. “*α*” and “*φ*” indicate the amplitude and the phase of the fitted cosine curve. (B) Histograms of amplitude (*α*) of expressed genes. Left: *Ahg*. Right: *Ath*. The regions with *α* > 1 are colored. (C) Histograms of phase (*φ*) of expressed genes. Left: *Ahg*. Right: *Ath*. The regions with *α* > 1 are colored. (D) Venn diagram of *Ahg* and *Ath* SO genes. Left circle: *Ahg*. Right circle: *Ath*. (E) Scatter plot of amplitudes in *Ahg* and *Ath*. Only genes with *α* > 1 in both species are colored according to density using the *densCols* function in R. (F) Scatter plot of phases in *Ahg* and *Ath*. Only genes with *α* > 1 in both species are colored according to density using the *densCols* function in R. (G) Venn diagram of genes that passed amplitude and phase criteria. Left circle: genes that passed amplitude criteria (*Δα* < ± 1). Right circle: genes that passed phase criteria (*Δφ* < ± 45 days). (H) Seasonal expression plots of *LHY*. From left, fitted cosine curve plot of *Ahg* and *Ath*, observed value and fitted line plot of *Ahg* and *Ath*, respectively.

Based on this criterion, 4312 and 4016 genes were identified as SO genes in *Ahg* and *Ath*, respectively. Although phases of these SO genes were concentrated in summer (June to August) or winter (December to February) in both *Ahg* and *Ath*, the phase of *Ahg* SO genes was wider than that of *Ath* SO genes (Fig. **3C**). In *Ahg*, 48% (2099 genes) and 33.5% (1445 genes) of the 4312 SO genes had phases distributed in summer and winter, respectively. In *Ath*, 52% (2125 genes) and 42% (1693 genes) of the 4016 SO genes had phases distributed in summer and winter, respectively. Among 4312 *Ahg* SO genes and 4016 *Ath* SO genes, 1757 genes overlapped (Fig. **3D**). By comparing the amplitude and phase in *Ath* and *Ahg*, we defined genes whose amplitude difference was < ± 1 and whose phase difference was < ± 45 days as seasonality reproduced genes (SR genes) (Fig. **3E-G**). Between *Ahg* plants grown in the field and *Ath* plants cultured in the SGCmini for 7 d, 946 genes were found to be SR genes.

To elucidate the functional aspects of the SR genes, we tested the enrichment of genes with a functional annotation. We found that 63 GOs were significantly enriched (adjusted *P* < 0.05, Supplementary Table **S3**). For example, genes with annotations of ‘response to temperature stimulus’ (GO: 0009266, adjusted *P* = 0.009) were enriched. One of these genes, *AhgLHY* (*LATE ELONGATED HYPOCOTYL 1*), which is a core clock gene, has been reported in a previous study as a gene that showed seasonally oscillating expression at noon. *AthLHY* (AT1G01060.1) had almost the same seasonal trend as *AhgLHY* (Fig. **3H**). It has been reported that the amplitude of the circadian rhythm of *AhgLHY* changes seasonally, and that sampling at noon results in a seasonal trend of high expression in winter and low expression in summer (Nagano et al., 2019).

### Several SO genes required extreme high temperatures to reproduce their seasonal trends

Several of the genes annotated as “response to temperature stimuli” were SR genes, such as *LHY*, and others were not. One possible reason for the non-mimicry is the mild temperature setting of the SGCmini. For example, although we chose the average temperatures, in summer, plants transiently face higher temperatures in the field. To mimic this condition, a high-temperature experiment was performed. In the experiment, daytime temperature in the August condition was changed to 32 °C or 35 °C (Fig. **4A,B**).

**Fig. 4.**
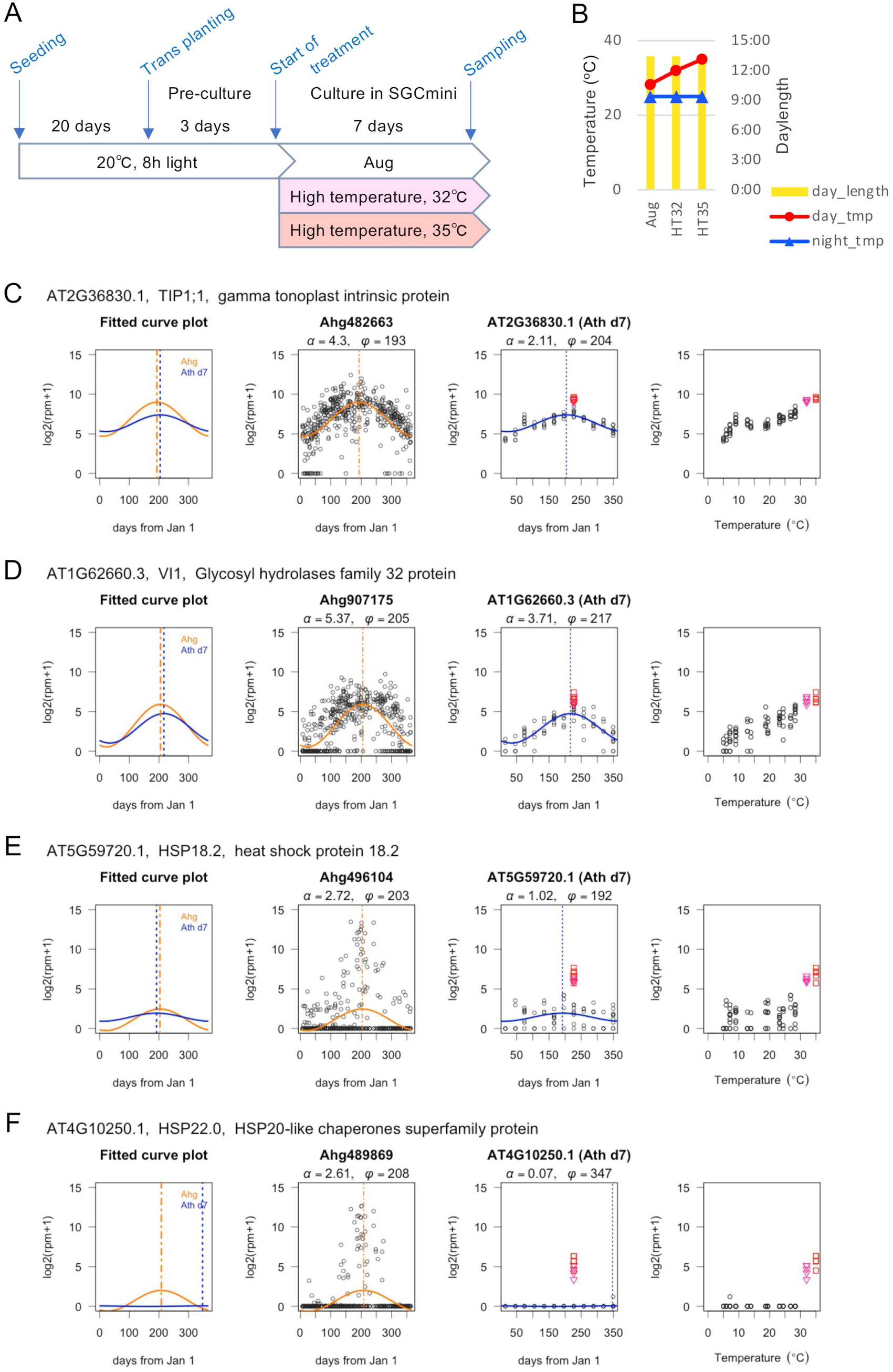
Effect of higher temperature on gene expression. (A) Schematic diagram of culture conditions. (B) Conditions of the high temperature experiment. Yellow bars: daylength; red circles: daytime temperatures; blue triangles: nighttime temperatures. (C) Gene expression plot of *TIP1;1*. From left, fitted cosine curve plot of *Arabidopsis halleri* ssp. *gemmifera* (*Ahg*) and *Arabidopsis thaliana* (*Ath*), observed value and fitted line plot of *Ahg* and *Ath*, respectively, and a plot of temperature and expression levels. Black circles indicate observed values. Orange (*Ahg*) and blue (*Ath*) lines show the fitted cosine curve. Orange and blue dashed lines indicate the phases of *Ahg* and *Ath*, respectively. Pink triangles and red squares indicate observed values when plants were grown at 32 °C and 35 °C. (D) *VI1*. (E) *HSP18.2*. (F) *HSP22.0*.

*GAMMA TONOPLAST INTRINSIC PROTEIN 1* (*TIP1;1*, AT2G36830) and *VACUOLAR INVERTASE 1* (*VI1*, AT1G62660) showed pseudo-seasonal oscillations in *Ath* (Fig. **4C,D**). Although their phases were almost the same as that of *Ahg*, their amplitudes were smaller than the threshold (amplitude difference < 1). When cultivated under higher temperatures, the expressions of these genes were successfully elevated to a level similar to that in *Ahg*.

In contrast, in the field, *Ahg* orthologs of *HEAT SHOCK PROTEIN 18.2* (*HSP18.2*, AT5G59720) and *HSP22* (AT4G10250) showed heterogeneous high gene expression levels among plant individuals in summer (Fig. **4E,F**). Heterogeneous expression is speculated to be a response to the heterogeneous microenvironment around individual plants, that is, locally elevated temperatures (Nagano et al., 2019). Although the expressions of *HSP18.2* and *HSP22* were not high under August conditions (daytime temperature: 28.3 °C), these expressions were notably elevated at 32 °C and 35 °C (Fig. **4E,F**). Thus, for some genes, non-average conditions, such as the highest or lowest temperatures in a day, are necessary to mimic the seasonal trends in gene expression.

### Shorter cultivation periods resulted in smaller morphological seasonal changes

To investigate the effect of culture period length, plants were cultured in the SGCmini for 1 or 3 d under the condition of each month (Fig. **5**; Supplementary Fig. **S3** and **S4**). As shown in Fig. **5A**, pre-culture periods were changed depending on the culture period in each month condition to make the days after sowing uniform when morphology was measured. As the culture period in each month became shorter, the pseudo-seasonal change in morphology became more moderate (Fig. **5B,C**). In the 3-d culture, total leaf area increased under July and August conditions (Fig. **5B**; Supplementary Fig. **S3D**). Petiole length increased under warmer conditions (June to September). The number of leaves slightly increased under the July, August, and September conditions (Fig. **5B**; Supplementary Fig. **S3F**). The leaf blade length slightly increased under the July condition (Fig. **5B**; Supplementary Fig. **S3G**). In the 1-d culture, there was no significant difference in plant growth (Fig. **5B**; Supplementary Fig. **S4**). Morphological observations showed that the differences among the conditions of each month were smaller in the shorter culture period lengths.

**Fig. 5.**
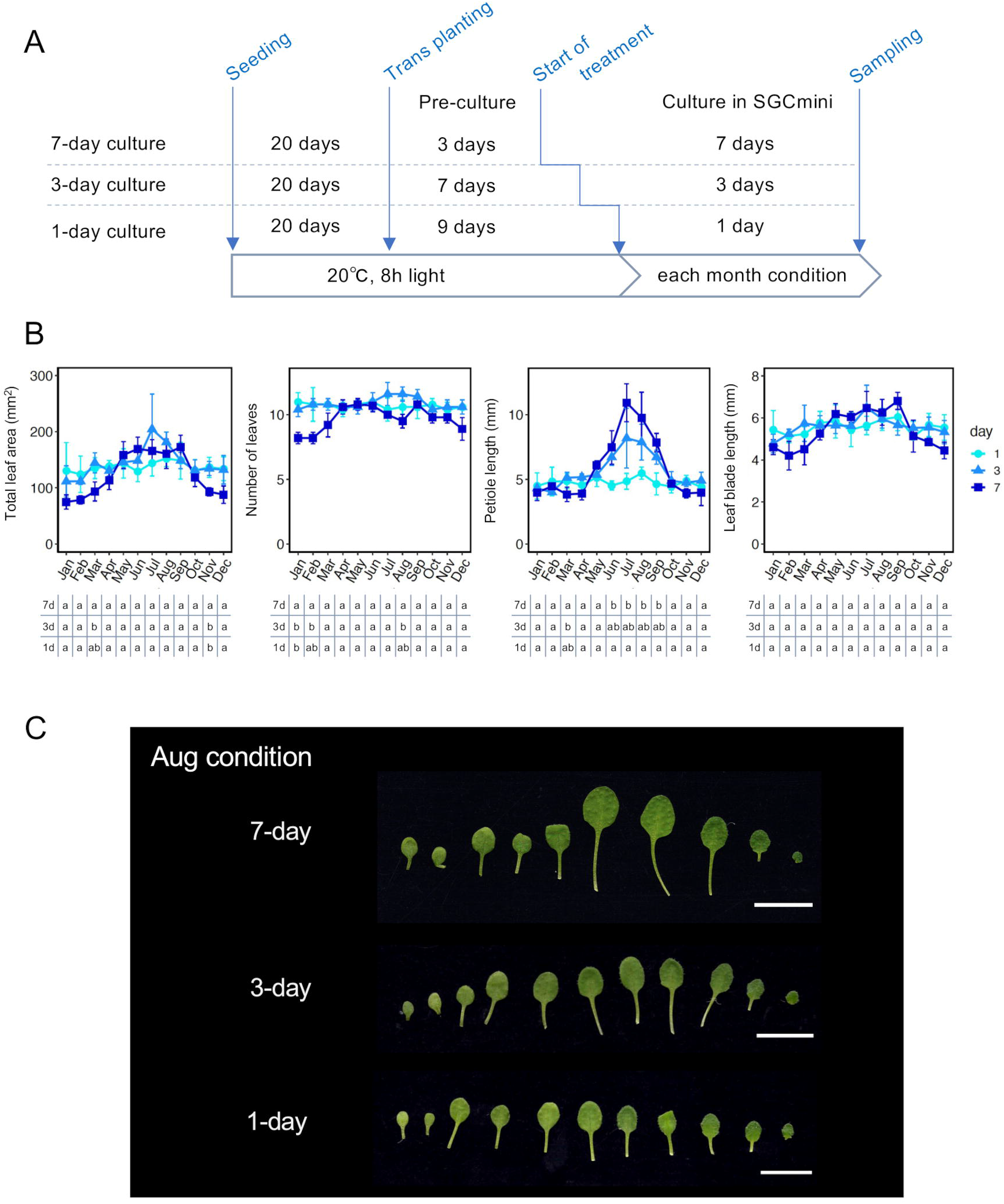
Comparison of morphological traits of *Arabidopsis thaliana* (*Ath*) plants grown in the SGCmini for 1, 3, or 7 d. (A) Schematic diagram of culture conditions. (B) Morphological traits. From left, total leaf area, number of leaves, maximum petiole length, and maximum leaf blade length. Dark blue squares, middle blue triangles, and light blue circles indicate data for plants grown in the SGCmini for 7, 3, and 1 d, respectively. n = 4–9. Different letters below the plots indicate differences between culture periods at *P* < 0.001 using ANOVA and Tukey’s HSD test. (C) Leaves of plants grown under the August condition. From the top, 7-, 3- and 1-d cultures. The youngest leaves are on the right. Bar = 10 mm.

Differences between culture periods (7, 3, and 1-d) at *P* < 0.001 using ANOVA and Tukey’s HSD test are shown in Fig. **5B**. Taking the number of tests into account, we chose a conservative threshold (results of differences at *P* < 0.05 are shown in Supplementary Tables **S4–7**). Significant differences in total leaf area were detected between 7- and 3-d cultures in March and between 7- and both 3- and 1 d cultures in November. Significant differences in the number of leaves were detected between 7- and both 3- and 1-d cultures in January and between 7- and 3-d cultures in February and August, indicating that development of visible leaves tended to be suppressed under these conditions in the 7-d culture. Significant differences in maximum petiole length were detected between 7- and 1-d cultures from June to September, indicating that petiole elongation tended to be promoted under these conditions as the culture period increased. A significant difference in maximum petiole length was also detected between the 7- and 3-d cultures in March. No significant difference was detected in the leaf blade length between the culture periods.

### Each seasonally oscillating gene required a different length of cultivation period to reproduce its seasonal trend

RNA-Seq was performed to clarify how the length of the culture period affected the pseudo-seasonal trend of gene expression in *Ath*. The SO genes and SR genes were defined in the same manner as the analysis of 7-d culture data (Fig. **6**; Supplementary Fig. **S5** and **S6**). The number of SO genes was similar in the 7-d culture (4016 genes) and 3-d culture (3975 genes) (Fig. **6A**). The number of SO genes in the 1-d culture was greater than that in the 3- and 7-d cultures (4575 genes). The number of SO genes that were annotated as orthologs of *Ahg* SO genes showed the same tendency (7-d culture: 1757 genes, 3-d culture: 1766 genes, 1-d culture: 1886 genes). As the culture period became shorter, the phase distribution became more concentrated in summer and winter (Fig. **6B**). A total of 412 SR genes were shared with three conditions (Fig. **6C**). Examples of the shared genes were *LHY* and *SUCROSE SYNTHASE 1* (*SUS1*, AT5G20830). These genes showed almost the same seasonal trends regardless of the length of the culture period (Fig. **7A,B**).

**Fig. 6.**
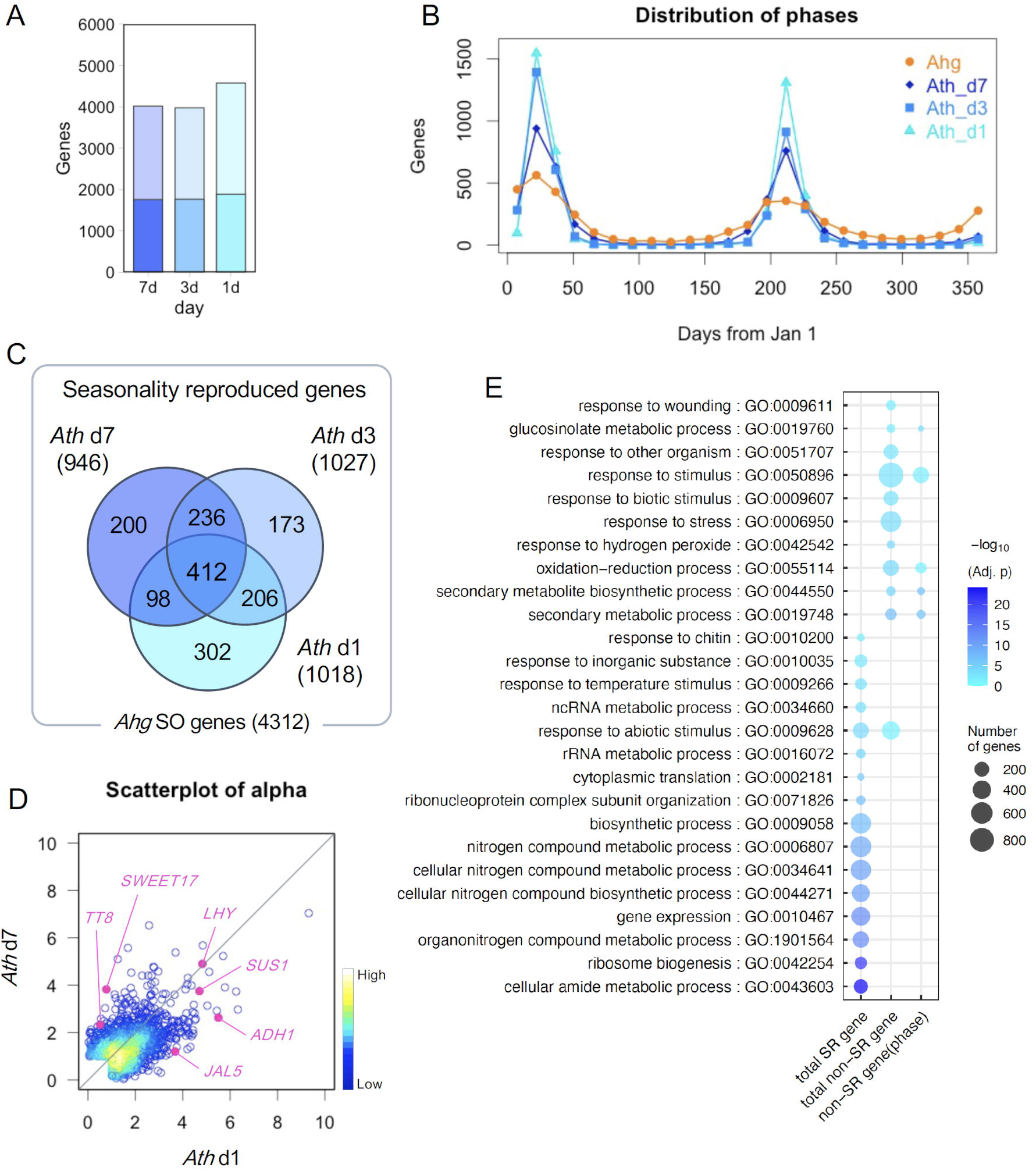
Comparison of seasonal trends of gene expression between different culture periods. (A) Comparison of the number of seasonally oscillating (SO) genes. Darker regions indicate the number of *Arabidopsis thaliana* (*Ath*) SO genes common with those of *Arabidopsis halleri* ssp. *gemmifera* (*Ahg*) defined in the field experiment data. The x-axis indicates the number of days of the length of the incubation period in SGCmini. (B) Comparison of the distribution of phases of SO genes. Orange circles: *Ahg*; dark blue rhombuses: *Ath* 7-d culture; blue squares: *Ath* 3-d culture; light blue triangles: *Ath* 1-d culture. (C) Venn diagram of seasonality reproduced genes (SR genes). Left circle: *Ath* 7-d culture; right circle: *Ath* 3-d culture; bottom circle: *Ath* 1-d culture. (D) Scatter plot of the alpha of SR genes in the 7-d culture (vertical axis) and 1-d culture (horizontal axis). Each point is colored based on the local density of points. Pink circles indicate genes shown in Fig. **7**. (E) Enriched biological process gene ontology (GO) terms. The “total SR gene” indicates the total SR number of genes that mimicked the seasonal trend of *Ahg* in at least one of the three conditions (7-, 3-, and 1-d culture) and corresponds to the colored region in Fig. 6C. The “total non-SR gene” indicates non-SR genes that did not mimic the seasonal trend of *Ahg* and corresponds to the non-colored region in Fig. 6C. Among the “total non-SR genes,” “non-SR gene (phase)” indicates genes that passed the amplitude criteria but were not in phase with *Ahg*. Circle size indicates the number of genes. Color scale indicates the adjusted *P* values in the GO enrichment analysis.

**Fig. 7.**
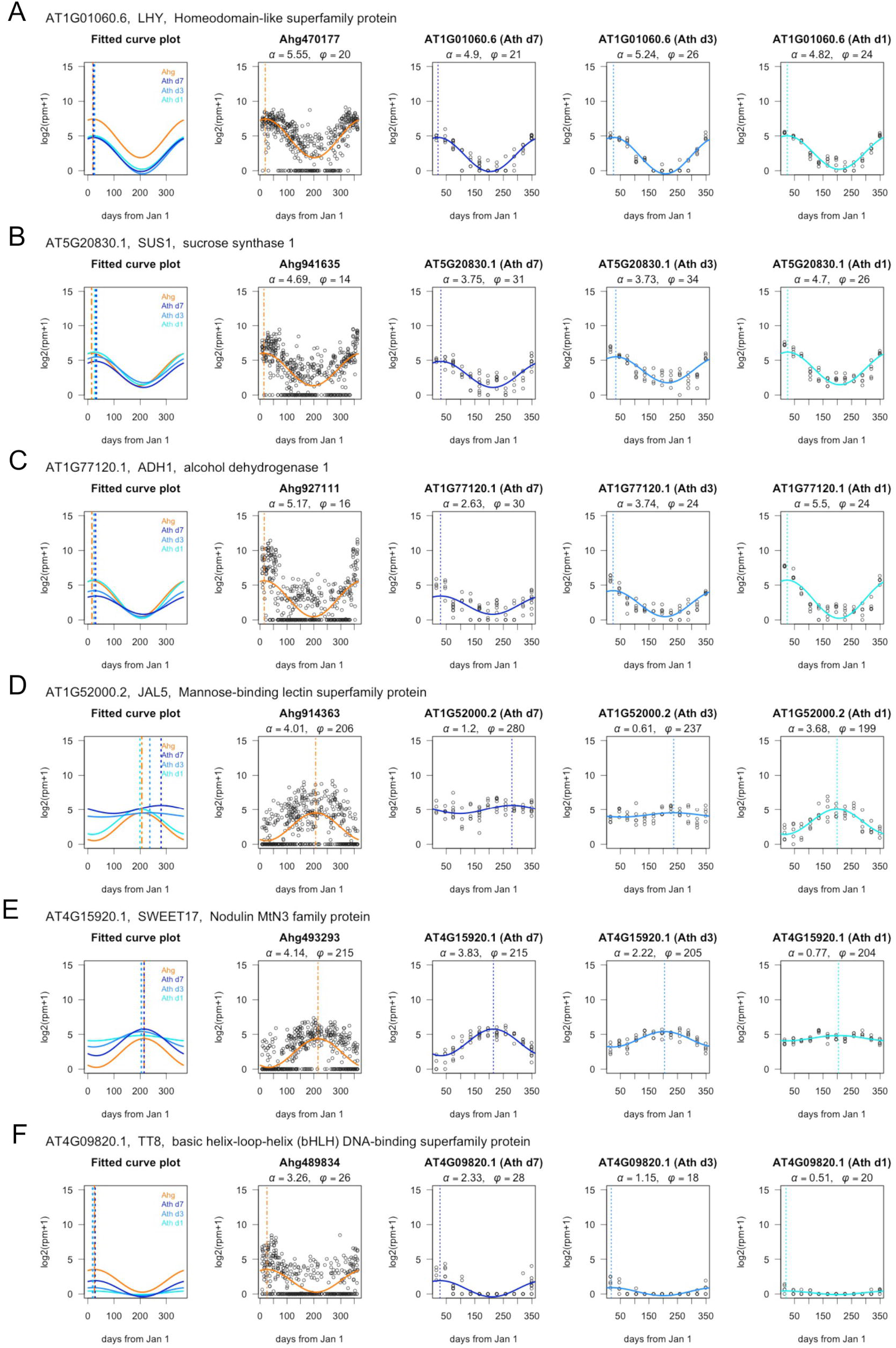
Seasonal expression plots of seasonality reproduced genes. From left, fitted cosine curve plot of *Arabidopsis halleri* ssp. *gemmifera* (*Ahg*) and *Arabidopsis thaliana* (*Ath*), observed value and fitted line plot of *Ahg* and *Ath* in the 7-, 3-, and 1-d cultures, respectively. (A) *LHY*. (B) *SUS1*. (C) *ADH1*. (D) *JAL5*. (E) *SWEET17*. (F) *TT8*.

Interestingly, hundreds of genes mimicked the *Ahg* seasonal trend only when the culture period was longer or shorter (SR genes in 7-d culture but not in 1-d culture: 436 genes, SR genes in 1-d culture but not in 7-d culture: 508 genes). *ALCOHOL DEHYDROGENASE 1* (*ADH1*, AT1G77120) and *JACALIN-RELATED LECTINS 5 (JAL5*, AT1G52000) were classified as SR genes in the 1-d culture but not in the 7-d culture (Fig. **6D**, **7C,D**). As the culture period increased, the amplitude of these genes decreased. *SWEET17* (AT4G15920) and *TRANSPARENT TESTA 8* (*TT8*, AT4G09820) were classified as SR genes in the 7-d culture but not in the 1-d culture (Fig. **6C**, **7E,F**). The amplitude of these genes decreased as the culture period became shorter. TT8 is a bHLH transcription factor that regulates flavonoid biosynthesis with other factors (Li, 2014; Xu et al., 2015). Anthocyanin regulatory and biosynthetic pathways and predicted seasonal expression plots of *Ahg* and *Ath* are shown in Supplementary Fig. **S7–10**. Among these genes, the amplitudes of the pseudo-seasonal trend of *DIHYDROFLAVONOL 4-REDUCTASE* (*DFR*, AT5G42800), *LEUCOANTHOCYANIDIN DIOXYGENASE* (*LDOX*, AT5G42800), *PRODUCTION OF ANTHOCYANIN PIGMENT 1* (*PAP1*, AT1G56650), and *PAP2* (AT1G66390) became larger as the culture period increased, similar to *TT8* (Supplementary Fig. **S7A,C, S8**, and **S10**). In our criteria, *F3’H, MYB11, MYB111, PAP1, PAP2, TT8, GL3*, and *EGL3* were defined as SO genes in *Ahg*. Among them, only *TT8* and *PAP2* were defined as SR genes in *Ath*.

The reason for the difference in the amplitude of the seasonal trend among the three conditions in *Ath* could be the difference in the length of the culture period (7, 3, and 1 d) and/or the difference in the plant age at the start of the treatment in the SGCmini (23, 27, and 29 d). To investigate the effect of plant age on the amplitude of SO genes, plants at 23 d after seeding were cultured in the SGCmini for 1 d under the January and August conditions (Supplementary Fig. **S11A**), and RNA-Seq was performed. The expression levels of SO genes of younger plants tended to have slightly smaller differences between the January and August conditions (Supplementary Fig. **S11B**). However, the variation due to the difference in the length of the culture period was greater than that due to plant age (Supplementary Fig. **S11B**–**D**).

In total, 1627 genes were identified as SR genes in at least one of the three conditions (Fig. **6C**; Supplementary Fig. **S2B,** Table **S8**). To elucidate the functional aspects of the SR and non-SR genes, we performed GO enrichment analysis (adjusted *P* < 0.05, Supplementary Tables **S9-11**). In all SR genes, the GO terms “gene expression,” “response to abiotic stimulus,” and “response to temperature stimulus” were enriched (Fig. **6E**). The GO term “response to abiotic stimulus” was enriched only in the SR gene in 7-d culture (Fig. **S12**, Supplementary Tables **S12-14**). In all non-SR genes, the GO terms “secondary metabolic process,” “response to biotic stimulus,” and “response to abiotic stimulus” were enriched. The GO terms “secondary metabolic process” and “glucosinolate metabolic process” were enriched in genes that passed the amplitude criteria but were not in phase with *Ahg*. This indicates that several SO genes associated with these GO terms require other factors (other environmental stimuli, etc.) to mimic the seasonal trend of *Ahg* in the appropriate phase, even though they were induced under mimic conditions in the present experiment.

## Discussion

Short-term incubation of *Ath* plants in the SGCmini under conditions representing each month could mimic the seasonal trends of 1627 SO genes in *Ahg*. The results of the 7-, 3-, and 1-d culture periods revealed that the culture period required for mimicry differed depending on the SO gene (Fig. **6C**, **7;** Supplementary Fig. **S8**). For some SO genes, the length of the culture period seems to be unrelated to the mimicking of seasonal trends of *Ahg* gene expression in the field because 412 SR genes were observed under all three conditions. These SO genes may ensure robustness to environmental changes by responding accurately to the current environment, regardless of the past environment. *LHY* is one of these SO genes (Fig. **7A**); it is known as a core clock gene that contributes to temperature compensation of the circadian clock (Gil and Park, 2019; Gould et al., 2006; Salomé et al., 2010). In field environments, temperatures can change drastically, even within a few days. The constant responsiveness of *LHY* may contribute to maintaining the circadian rhythm under fluctuating environmental conditions of the field. In our previous study, we found that 1792 SO genes had both seasonal and diurnal oscillations (62.2% of SO genes)(Nagano et al., 2019). Recently, the complex effects of clock genes in the natural environment have received increasing attention (Oravec and Greenham, 2022) because rising temperatures due to global warming disrupt existing plant responses to day length and temperature during seasonal events. Accordingly, we have now established a convenient experimental system using SGCmini. We will be able to assess the contribution of each gene (such as clock genes) to the seasonal transcriptome from pseudo-seasonal experiments using mutants of each gene in *Ath*.

Conversely, there were several SO genes that mimicked the *Ahg* seasonal trend only when the culture period was shorter (Fig. **7C,D**). *ADH1* was one of these SO genes. Song *et al*. showed that *ADH1* transcription was induced at 3 h following 4 °C stress treatment, reached the highest expression level at 24 h, then declined (Song et al., 2017). Since its amplitude decreased under longer incubations, *ADH1* seems to respond to environmental changes (in this experiment, changes in temperature and daylength from pre-culture conditions to conditions of each month in the SGCmini), rather than to absolute values at the time, as LHY does. *ADH1* is not only involved in the cold stress response (Song et al., 2017), but its transcription is also known to be induced by various other stresses (Shi et al., 2017; Winter et al., 2007). SO genes such as *ADH1*, which mimicked field seasonal trends only in shorter culture periods, may play a role in responding to rapid environmental changes (within 1 day) due to their transient expression.

Several SO genes mimicked field seasonal trends only in longer culture periods, such as *SWEET17* and *TT8* (Fig. **7E,F**). In *Ath*, late biosynthetic genes (LBGs) of the anthocyanin biosynthetic pathway, *DFR* and *LDOX*, and their transcriptional regulators, *TT8, PAP1*, and *PAP2*, showed an increase in the amplitude of the seasonal trend with the extension of the culture period, suggesting a regulatory mechanism that responds to continuous environmental stimuli such as low temperature (Supplementary Fig. **S7–10**). One possible mechanism driving this is epigenetic regulation. Recent studies have revealed that epigenetic regulation is involved in the regulation of anthocyanin biosynthesis gene expression (Cai et al., 2019; Zheng et al., 2019). In the field, analysis of seasonal histone modifications of *Ahg* suggests that histone H3 lysine 27 trimethylation is involved in gene expressions that depend on memory of longer-term (more than a few weeks) environmental conditions (Nishio et al., 2020). In our study, although the plants only experienced the treatment once, the memory of past intermittent stimuli, such as hot or cold temperatures, can also affect plant seasonal response (Leuendorf et al., 2020; Yamaguchi et al., 2021). Evaluating the contribution of longer-term (month-scale) and intermittent environmental stimulus inputs to seasonal trends is one of the next challenges in the study of SO genes.

The differences in the temporal responsiveness of SO genes may contribute to the variation in the phase of the seasonal trend (Fig. **6B**). In addition, integrated response to multiple stimulus information may also be responsible for the wide distribution of phases in field *Ahg*. The phase of pseudo-seasonal trends of *PAP1* in *Ath* did not match that in *Ahg* (Fig. **S7C**). In SGCmini, *PAP1* expression is up-regulated in winter conditions, but in the field, its expression is higher in summer than in winter. Although previous studies have shown that *PAP1* and *PAP2* are up-regulated at low temperatures (Petridis et al., 2016), *PAP2*, rather than *PAP1* is suggested to play a more specific role in the accumulation of flavonoids at cold temperatures (Hannah et al., 2006). *PAP1* expression is also induced by other stimuli, such as high light intensity (Li et al., 2016; Shi and Xie, 2010). An integrated response to multiple stimulus information may also be the reason for the enrichment of GO terms like “response to stress” or “response to abiotic stimuli” in non-SR genes (Fig.**6E**). For some SO genes, multiple stimulus inputs in the field may form seasonal trends that are not correlated with a single environmental stimulus, such as temperature.

Alternatively, multiple seasonal stimuli may mask seasonal oscillations. Early biosynthetic genes (EBGs) and LBGs, except *F3’H*, did not show seasonal oscillations in *Ahg* in the field (Supplementary Fig. **S7** and **S8**). However, their transcriptional regulators showed seasonal oscillations in various phases (Supplementary Fig. **S7, S9,** and **S10**), suggesting that the constitutive expression of EBGs and LBGs in the field is the outcome of the integration of multiple environmental stimuli by transcriptional regulators. Flavonoid biosynthesis is affected not only by low temperature, but also by light quality, drought, nutrition, and virus infection (Cai et al., 2019; Cominelli et al., 2008; Honjo et al., 2020; Lillo et al., 2008). While highly oscillating genes are eye-catching, it is also important to remember that seemingly non-oscillating genes may be involved in plant responses to seasonal changes. Such a complex response mechanism of secondary metabolites may be one of the reasons why the GO term “secondary metabolic process” was detected in the GO analysis of SO genes that could not be mimicked (Fig. **6E**).

In conclusion, we successfully mimicked the seasonal trends of approximately 38% of *Ahg* SO genes using *Ath* plants by incubating them for 1 to 7 days in an experimental system set at the average temperature and daylength of each month. The results indicate that the field seasonal trend of these genes is shaped by day-scale responses to environmental stimuli rather than month-scale responses to long-term inputs. Conversely, the seasonal trends of 62% of the SO genes could not be mimicked. Although one of the reasons could be the lack of long-term inputs of environmental stimuli, there are many other possibilities. For example, species differences between *Ahg* and *Ath* and environmental stimuli which were not replicated in the SGCmini (fluctuation of environmental stimuli, biotic stress, etc.) may also be responsible. We will next clarify which of these possibilities is important for seasonal trends of non-mimicked SO genes in *Ahg*. This is the first comprehensive study of the day-scale response of SO genes to environmental stimuli. In order to adapt to complex environmental changes in the field, plants likely interpret seasonal changes by converting environmental stimuli into gene expression at different temporal resolutions (minute-, day- and month-scale). Although the functions of many SO genes are still unknown, the results of this study and the experimental system can be useful for investigating these functions.

## Materials and Methods

### Smart growth chamber mini (SGCmini)

The system configuration of the developed smart growth chamber mini (SGCmini) is shown in Supplementary Fig. **S1A**. The proposed system consisted of an incubator body, Peltier unit, circuit unit, and control unit. Supplementary Fig. **S1B** depicts the proposed system, excluding the control unit. The incubator body was made of acrylic; the bottom surface was made of black acrylic and the rest was transparent acrylic (thickness: 10 mm). The transparent acrylic parts were all integrated and had holes into which the Peltier unit could be attached (described below). The outer dimensions of the transparent acrylic part were 80 mm × 200 mm × 200 mm.

The appearance of the Peltier unit is shown in Supplementary Fig. **S1C**. This unit comprised a 40 × 40 mm Peltier device, two heat sinks, and two fans. The heat sinks were adhered on both sides of the Peltier device and the black insulation sheets enclosed them. The side with the small heat sink was inserted through the hole in the incubator body and housed in the body. A small fan was installed to circulate the air inside the incubator. The side with the larger heat sink was for heat dissipation, which enabled air cooling with a large fan.

The circuit unit consisted of an Arduino Uno R3, a DC brush motor driver shield for Arduino (SHIELD-MD10, Cytron Technologies, Malaysia), and two temperature sensors (MCP9808, Microchip Technology, USA). Two temperature sensors were installed in the incubator. The circuit unit sends information from the temperature sensors to the control unit (PC) via serial communication.

The control unit was equipped with a GUI, which determined the extent of temperature control for heating and cooling to achieve the “target temperature set by the GUI” in the incubator. The proportional integral (PI) control determined the temperature control according to the current temperature in the incubator, and it was sent to the circuit unit as the control signal. According to the signal received, the circuit unit controlled the current sent to the Peltier unit by pulse-width modulation (PWM).

The GUI of the control unit is illustrated in Supplementary Fig. **S1D**. In this GUI, the temperature in the incubator could be set in increments of 0.5 °C. In addition to the current temperature, the temperature profile for the past 48 h could be displayed on a chart. The GUI also provided other functions, such as the output of a temperature control log and the capability to change the set temperature. The developed incubator could control the temperature with an error range of 0.2 °C within a range of −9 to +20 °C from the standard room temperature. For colder conditions, such as that for February, the SGCmini was operated in a growth chamber (LH-241SP, NK system Osaka, Japan) set below 15 °C.

For plant cultivation, the incubator body was painted black on four sides to block external light (Fig. **1A**). An LED light unit (25 pcs; color: red and blue; size: 30.8 × 30.8 cm; PPFD: approximately 35 μmol m^-2^ s^-1^) was placed on the top of the incubator body. The lighting of the LED light unit was controlled using a timer.

### Setting of culture condition in SGCmini

To mimic the seasonal expression of *Arabidopsis halleri* ssp. *gemmifera* (*Ahg*) in the field, simplified conditions of the environment of the *Ahg* sampling site (the Omoide River, Taka-cho, 35° 06’ N, 134° 55’ E, altitude 190–230 m above sea level) (Nagano et al., 2019) were set as culture conditions (Fig. **2A**). For daylength settings, sunrise and sunset times on the 15^th^ of each month in Nishiwaki city were obtained from the National Astronomical Observatory of Japan. For temperature settings, hourly average temperatures at Nishiwaki (averaged over 2006–2016) were obtained from the Japan Meteorological Agency. The average value of the temperature from sunrise to sunset was set as the daytime temperature, and the average value of the temperature from sunset to sunrise was set as the nighttime temperature. It was difficult to mimic temperatures below 5 °C, so when the temperature was below 5 °C, such as in January, the experiment was conducted with a 5 °C setting.

We also conducted a high-temperature culture experiment, in which the daytime temperature of the August condition was changed to 32 °C or 35 °C (Fig. **4A**).

### Plant materials

Seeds of *A. thaliana* (*Ath*) (Col-0, CS70000) were sown on 1/2 MS medium with 0.5% sucrose and 0.6% gellan gum in square Petri dishes and cultured at 20 °C with 8 h light for 20 d. Plants were transferred to 1/2 MS medium with 0.25% gellan gum in round plates and cultivated for 3, 7, or 9 d under the same conditions as the pre-culture to recover from the stress of transplanting. After the pre-culture, the round plates were placed in the SGCmini. Plants were cultivated under the conditions of each month for 7, 3, or 1 d (Fig. **2B** and **5B**). To equalize the age of the plants at the sampling date, we extended the pre-culture period to shorten the culture period. However, for a 1-day culture with short pre-culture conditions, the pre-culture period was set to 3 d in order to evaluate the effect of plant age at the start date of treatment (Supplementary Fig. **S12A**).

For morphometry, all leaves were cut off and scanned using a scanner (GT-S650, EPSON, Japan). Leaf area and leaf length were measured from the scanned images using ImageJ software. ANOVA and Tukey HSD were performed in R software to test for differences between conditions (*P* < 0.05).

### RNA-Seq

For RNA extraction, leaves larger than 1 mm without cotyledons and juvenile leaves were harvested from one plant as one sample between 11:00 and 13:00. The leaves were immediately frozen in liquid nitrogen and stored at −80 °C until extraction.

Samples were ground with two zirconia beads using a TissueLyser II (QIAGEN, MD, USA) with pre-chilled adapters at −80 °C. Total RNA from leaves was extracted using a Maxwell 16 LEV Plant RNA Kit (Promega, WI, USA). The amount of RNA was determined using Quant-iT RNA Assay Kit, broad range (Thermo Fisher Scientific, Waltham, MA, USA) and Tecan plate reader Infinite 200 PRO (Tecan, Männedorf, Switzerland). For RNA-Seq library preparation, 400 or 500 ng of total RNA per sample was used. The library was prepared using the Lasy-Seq v1.0 or v1.1 protocol (Kamitani et al., 2019) (https://sites.google.com/view/lasy-seq/). The quality of the library was assessed using a Bioanalyzer (Agilent Technologies). The libraries were sequenced with a HiSeq 2500 or Hiseq X (Illumina) to produce single reads of 50 bp (Library 1 and 2) or 150bp (Library 3).

Preprocessing and quality filtering of RNA-Seq data were performed using Trimmomatic-0.33 (Bolger et al., 2014). Preprocessed reads were mapped on transcript sequences in TAIR 10 with Bowtie1(v1.1.1) (Langmead et al., 2009) and quantified using RSEM-1.2.21(Li and Dewey, 2011). The output of the RSEM was analyzed using R (R Core Team, 2019).

RNA-Seq data of *A. halleri* were obtained from previous studies (Nagano et al., 2019). RNA-Seq data of *A. thaliana* and *A. halleri* were analyzed as follows. Samples with fewer than 10^5.5^ reads were excluded from the analysis. The attributes of the samples used in the analysis are listed in Supplementary Table **S1**. *Ahg* genes with an average log_2_(rpm+1) > 2 were designated as expressed genes (16158 genes) (Supplementary Fig. **S2**). A total of 90.3% of the expressed genes in *Ahg* (14587 genes) were annotated as orthologs of *Ath* genes by reciprocal BLAST. Subsequent seasonal expression analysis was performed using these annotated *Ahg*-expressed genes (14587 genes) and their orthologous genes in *Ath*.

### Analysis of seasonal expression

To compare the seasonal trends of *Ahg* and *Ath*, the expressions of the *Ahg*-expressed genes and their orthologous genes in *Ath* were fitted to a cosine curve using the *nls* function in R (Fig. **3A**). Amplitude and phase were obtained from each fitted cosine curve over a 1-y period. Genes with an amplitude > 1 were defined as seasonally oscillating genes (SO genes). Since the value of log_2_(rpm + 1) was fitted with a cosine curve, an amplitude > 1 indicates that the difference in seasonal oscillation is more than two-fold. We compared the amplitude and phase of *Ahg* and *Ath* genes to determine “seasonality reproduced genes (SR genes)” that could mimic the seasonal expression trends of *Ahg* in the field. A gene whose amplitude difference was smaller than 1 and whose phase difference was smaller than 45 d was defined as a SR gene.

### Gene ontology enrichment analysis

For gene ontology (GO) enrichment analysis, GO annotations for *Ath* genes were obtained from the TAIR database. Statistical tests of enrichment analysis were performed using the Fisher’s exact test function in R. A total of 14,472 expressed genes with GO annotations were included in the test. Multiple testing corrections were performed using FDR (Benjamini and Hochberg, 1995). Enriched GO terms are those with adjusted p <0.05. Representative GO terms were summarized based on the REVIGO output (http://revigo.irb.hr) (Supek et al., 2011) (allowed similarity = 0.5) and visualized with reference to a previous protocol (Bonnot et al., 2019).

## Data Availability Statement

The RNA-seq data were submitted to the NCBI Sequence Read Archive repository under the BioProject number PRJNA739047 (https://www.ncbi.nlm.nih.gov/bioproject/PRJNA739047). R scripts and data required for analysis (count data, etc.) are available via the GitHub repository (https://github.com/naganolab/AthSGCmini_pseudo-seasonal_RNA-Seq).

## Funding information

This work was supported by Japan Science and Technology Agency CREST [JPMJCR15O2], ACCEL [JPMJAC1403], FOREST [JPMJFR210B] and a project [JPNP18016] of the New Energy and Industrial Technology Development Organization (NEDO).

## Acknowledgements

We greatly appreciate Hiroshi Kudoh and his laboratory members for field data acquisition and discussions, Tetsuro Mimura for his advice and discussions on the experimental system, Kyoko Mogami for her help with RNA-Seq library preparation, and Motohiro Mihara and Hitoshi Ooshima of Dynacom Co., Ltd. (Chiba, Japan) for their support in RNA-Seq analysis, Daisuke Kyogoku for his advice on English language editing.

## Author Contribution

A.J.N. and Y.K. designed the research. H.T. and T.T. developed SGCmini. M. Kamitani, Y.H., M. Kashima, A.T. contributed to library preparation and data analysis. Y.K. performed experiments and data analysis. Y.K., H.T. and A.J.N. wrote the paper.

## Disclosures

### Conflicts of interest

No conflicts of interest declared.

## Notes

### Competing Interest Statement

The authors have declared no competing interest.

### Summary of Updates

We have changed the title of our paper to "Integration of short- and long-term responses to environmental stimuli shape seasonal transcriptome dynamics". We have added Fig .S12 and Supplementary Tables 12-14, and added several statements to the manuscript.

