## Supplementary material for "Integration of short- and long-term responses to environmental stimuli shape seasonal transcriptome dynamics": Fig. S

**The following Supporting Information is available for this article:**

**Fig. S1** Design of the smart growth chamber mini (SGCmini).

**Fig. S2** Summary of sample sets and gene sets in this study.

**Fig. S3** Morphological traits of *Arabidopsis thaliana* (*Ath*) plants grown in the SGCmini for 3 d.

**Fig. S4** Morphological traits of *Arabidopsis thaliana* (*Ath*) plants grown in the SGCmini for 1 d.

**Fig. S5** Comparison of seasonal trends of gene expression in *Arabidopsis halleri* ssp. *gemmifera* (*Ahg*) and *Arabidopsis thaliana* (*Ath*) grown in the SGCmini for 3 d.

**Fig. S6** Comparison of seasonal trends of gene expression in *Arabidopsis halleri* ssp. *gemmifera* (*Ahg*) and *Arabidopsis thaliana* (*Ath*) grown in the SGCmini for 1 d.

**Fig. S7** Anthocyanin regulatory and biosynthetic pathways and seasonal trend plots of *Arabidopsis halleri* ssp. *gemmifera* (*Ahg*) and *Arabidopsis thaliana* (*Ath*).

**Fig. S8** Seasonal expression plots of early biosynthetic genes (EBGs) and late biosynthetic genes (LBGs) in the anthocyanin biosynthetic pathway.

**Fig. S9** Seasonal expression plots of transcriptional regulators that regulate early biosynthetic genes (EBGs) in the anthocyanin biosynthetic pathway.

**Fig. S10** Seasonal expression plots of transcriptional regulators that regulate late biosynthetic genes (LBGs) in the anthocyanin biosynthetic pathway.

**Fig. S11** Comparison of gene expression patterns by length of pre-culture period.

**Fig. S12** Enriched biological process gene ontology (GO) terms in SR genes for each condition.

**The following Supporting Information Tables are available as a single Excel file, submitted separately:**

**Table S1** Attributes of the samples used in the present study

**Table S2** Setting of temperature and day length for each month in SGCmini.

**Table S3** The result of GO enrichment analysis of seasonality reproduced genes in the 7-day culture.

**Table S4** Test results for differences in total leaf area between culture periods.

**Table S5** Test results for differences in number of leaves between culture periods.

**Table S6** Test results for differences in max petiole length between culture periods.

**Table S7** Test results for differences in max leaf blade length between culture periods.

**Table S8** List of genes that could be mimicked Ahg seasonal trend.

**Table S9** The result of GO enrichment analysis of total SR genes that mimicked the seasonal trend of Ahg in at least one of the three conditions.

**Table S10** The result of GO enrichment analysis of non-SR genes that did not mimic the seasonal trend of Ahg.

**Table S11** The result of GO enrichment analysis of genes that passed the amplitude criteria but were not in phase with Ahg among the non-SR genes.

**Table S12** The result of GO enrichment analysis of SR genes in 7-d culture.

**Table S13** The result of GO enrichment analysis of SR genes in 3-d culture.

**Table S14** The result of GO enrichment analysis of SR genes in 1-d culture.

**Fig. S1** Design of the smart growth chamber mini (SGCmini). (A) Configuration of the proposed system. (B) Proposed system. (C) Peltier unit. (D) GUI provided by the control unit.

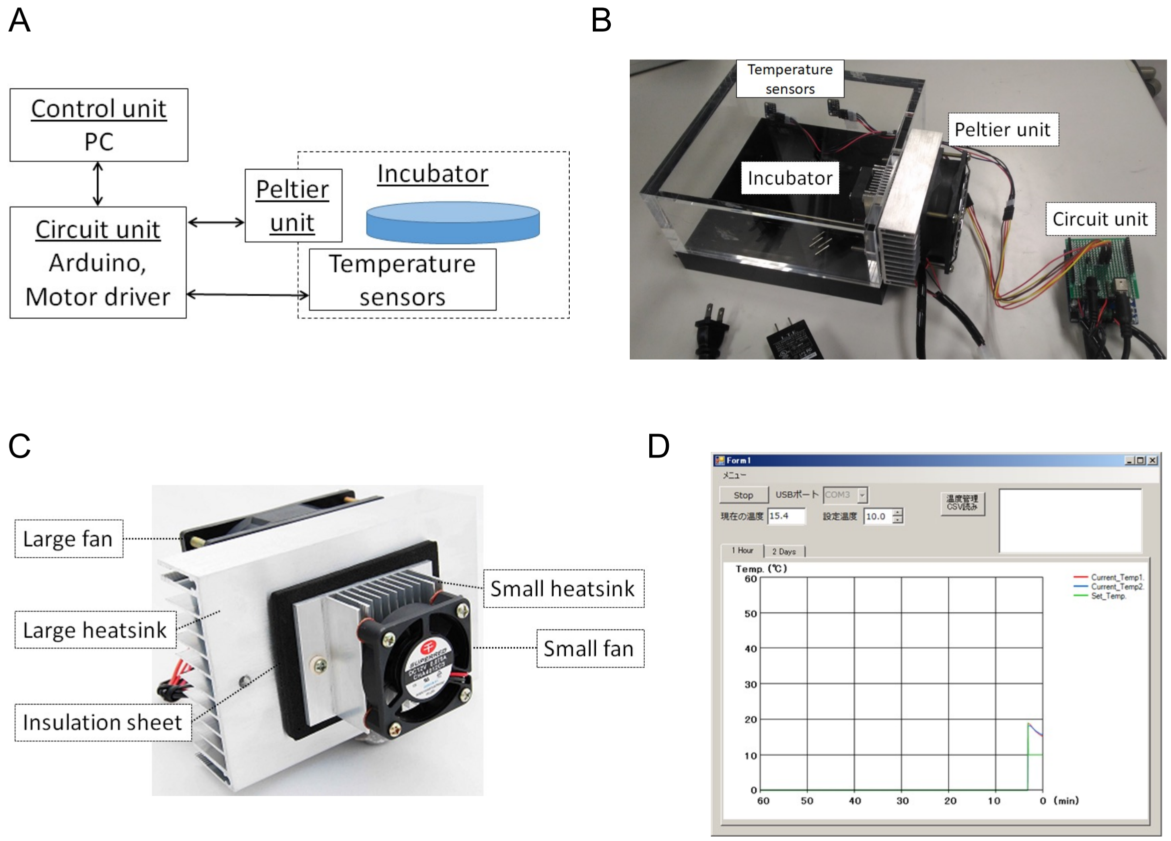

**Fig. S2** Summary of sample sets and gene sets in this study. (A) Sample sets in this study. (B) Summary of gene sets in this study. Histogram of mean expressions is shown in the inset. Pink line in the inset indicates the threshold of the expressed gene. "$\alpha$" and "$\varphi$" indicate the amplitude and the phase of fitted cosine curves.

A

| Species | *A. thaliana* | | | | | *A. halleri* |
|---|---|---|---|---|---|---|
| Type | Seasonal | | | | HT | Seasonal |
| name of treatment | d7 | d3 | d1 | s1 | HT | - |
| Days of treatment | 7 | 3 | 1 | 1 | 7 | - |
| Number of analyzed samples | 72 | 59 | 58 | 20 | 10 | 474 |

B

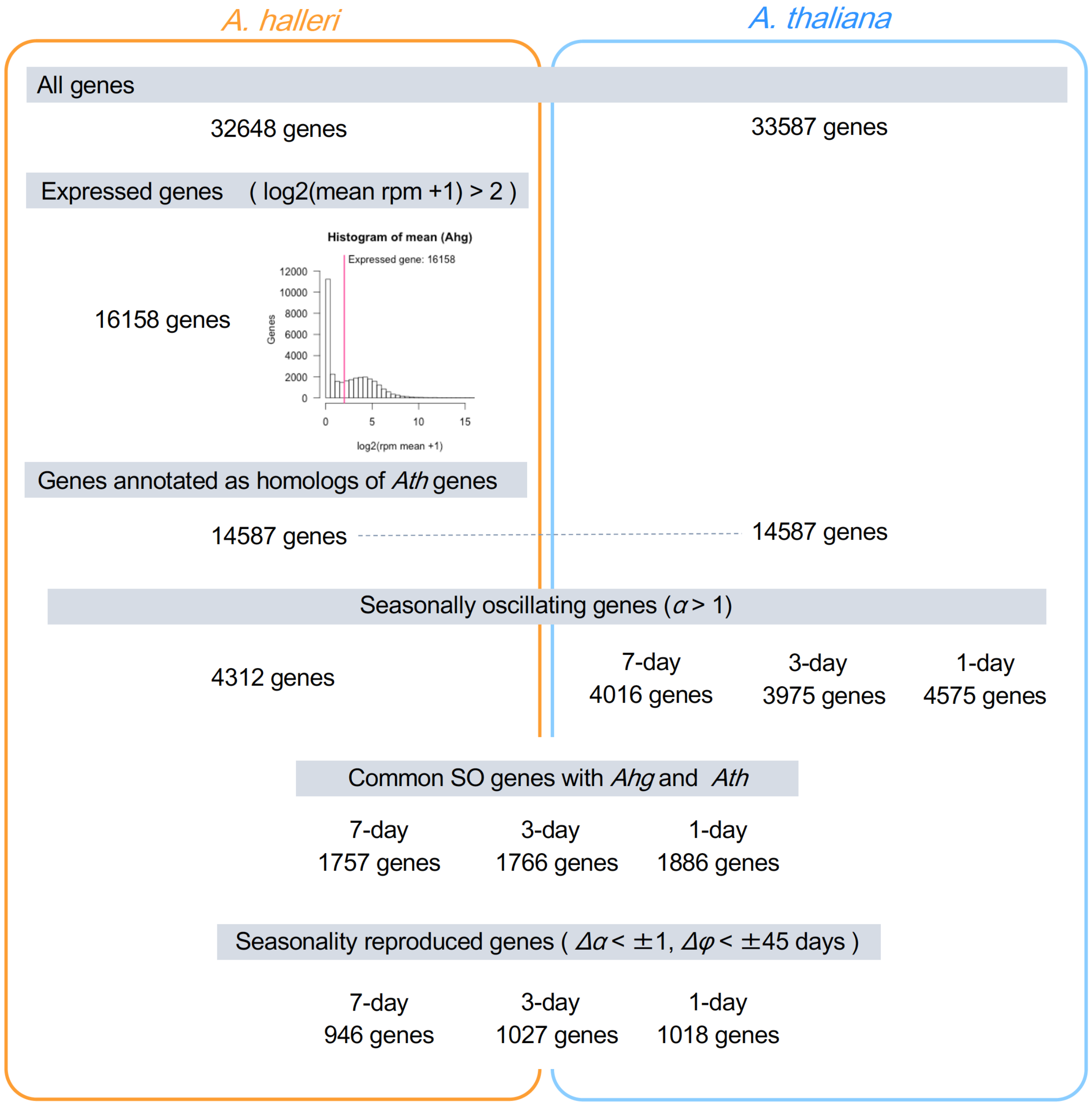

**Fig. S3** Morphological traits of *Arabidopsis thaliana* (*Ath*) plants grown in the SGCmini for 3 d. (A) Schematic diagram of the 3-d culture. Changes from the 7-d culture condition are shown in red. (B) Whole images of plants grown under February, May, and July conditions. (C) Leaves of plants grown under February, May, and July conditions. The youngest leaves are on the right. Bar = 10 mm. (D) Total leaf area. (E) Number of leaves. (F) Maximum petiole length. (G) Maximum leaf blade length. Bar: SD. n = 4–5. Different letters indicate differences between conditions at $P < 0.05$ using ANOVA and Tukey's HSD test.

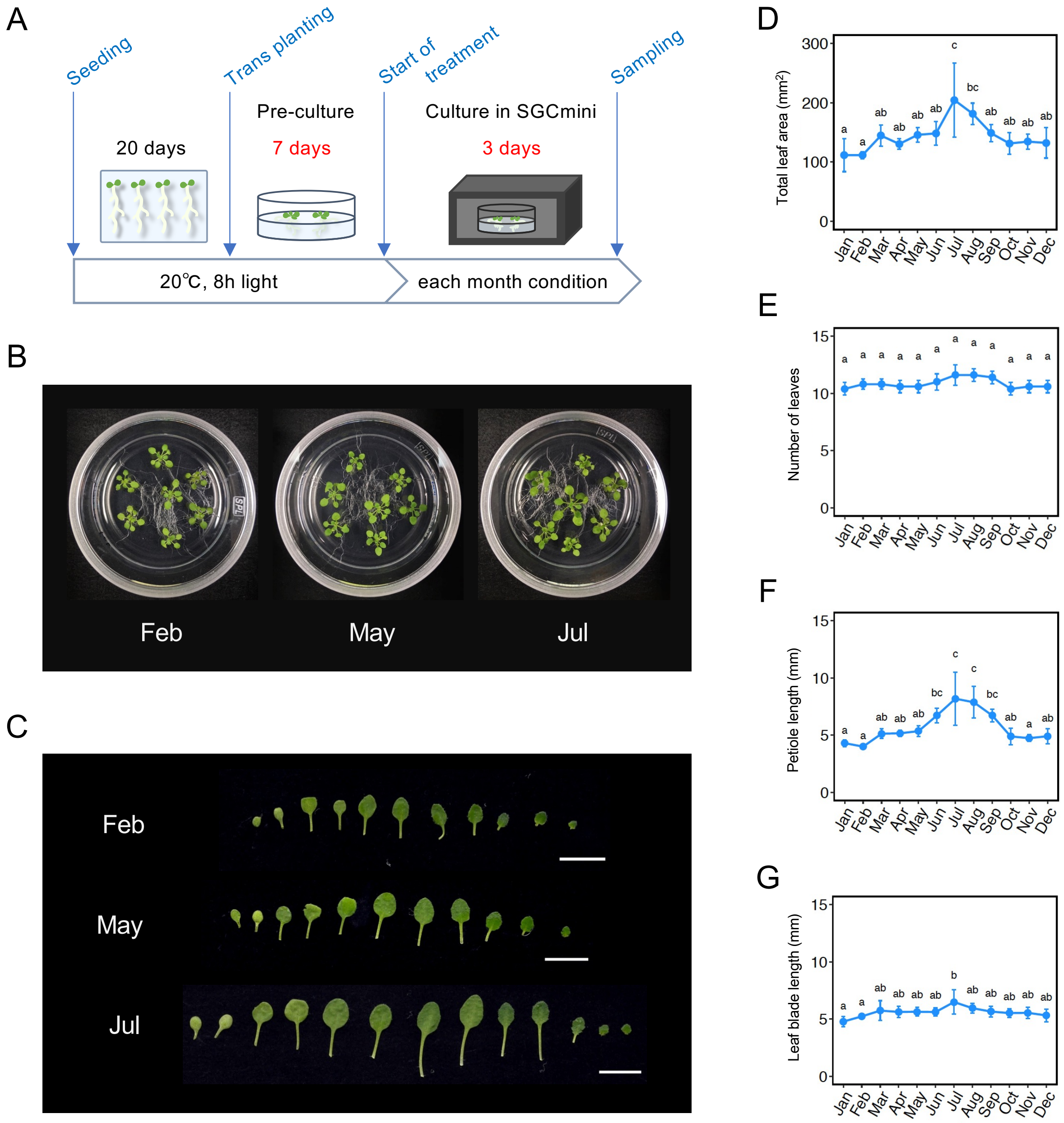

**Fig. S4** Morphological traits of *Arabidopsis thaliana* (*Ath*) plants grown in the SGCmini for 1 d. (A) Schematic diagram of the 1-d culture. Changes from the 7-d culture condition are shown in red. (B) Whole images of plants grown under February, May, and July conditions. (C) Leaves of plants grown under February, May, and July conditions. The youngest leaves are on the right. Bar = 10 mm. (D) Total leaf area. (E) Number of leaves. (F) Maximum petiole length. (G) Maximum leaf blade length. Bar: SD. n = 4–5. Different letters indicate differences between conditions at $P < 0.05$ using ANOVA and Tukey's HSD test.

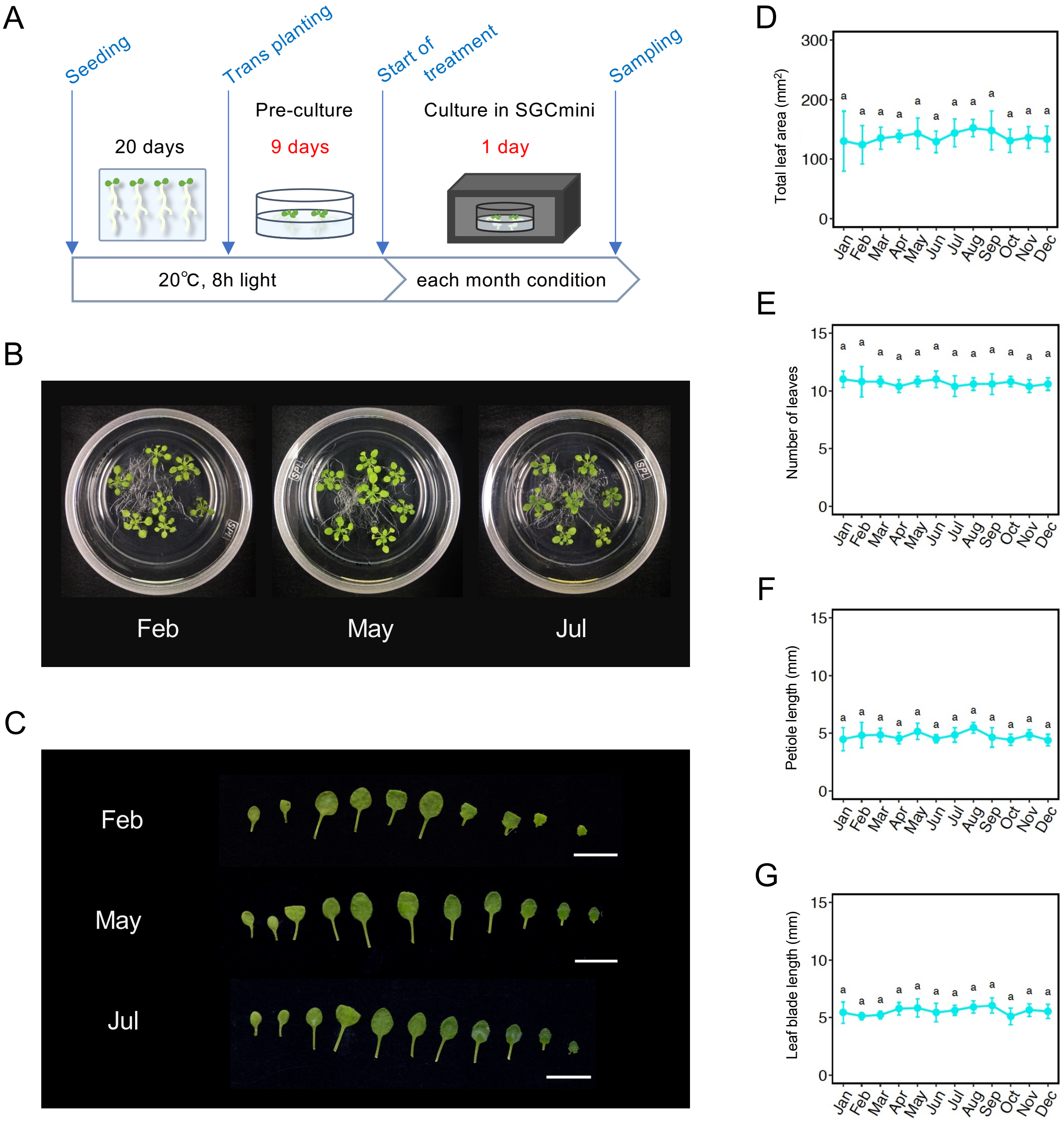

**Fig. S5** Comparison of seasonal trends of gene expression in *Arabidopsis halleri* ssp. *gemmifera* (*Ahg*) and *Arabidopsis thaliana* (*Ath*) grown in the SGCmini for 3 d. "α" and "φ" indicate the amplitude and the phase of fitted cosine curves. (A) Histograms of the amplitude of expressed genes. Left: *Ahg*. Right: *Ath*. (B) Histograms of the phase of expressed genes. Left: *Ahg*. Right: *Ath*. (C) Scatter plot of the amplitude of *Ahg* (vertical axis) and *Ath* (horizontal axis). Only genes with amplitude > 1 in both species were colored by the local density of points. (D) Scatter plot of the phase of *Ahg* and *Ath*. Coloring conditions are the same as (C). (E) Venn diagram of *Ahg* and *Ath* seasonally oscillating (SO) genes. Left circle: *Ahg*. Right circle: *Ath*. (F) Venn diagram of genes that passed the amplitude and phase criteria. Left circle: genes that passed the amplitude criteria ($Δα < 1$). Right circle: genes that passed the phase criteria ($Δφ < 45$ d).

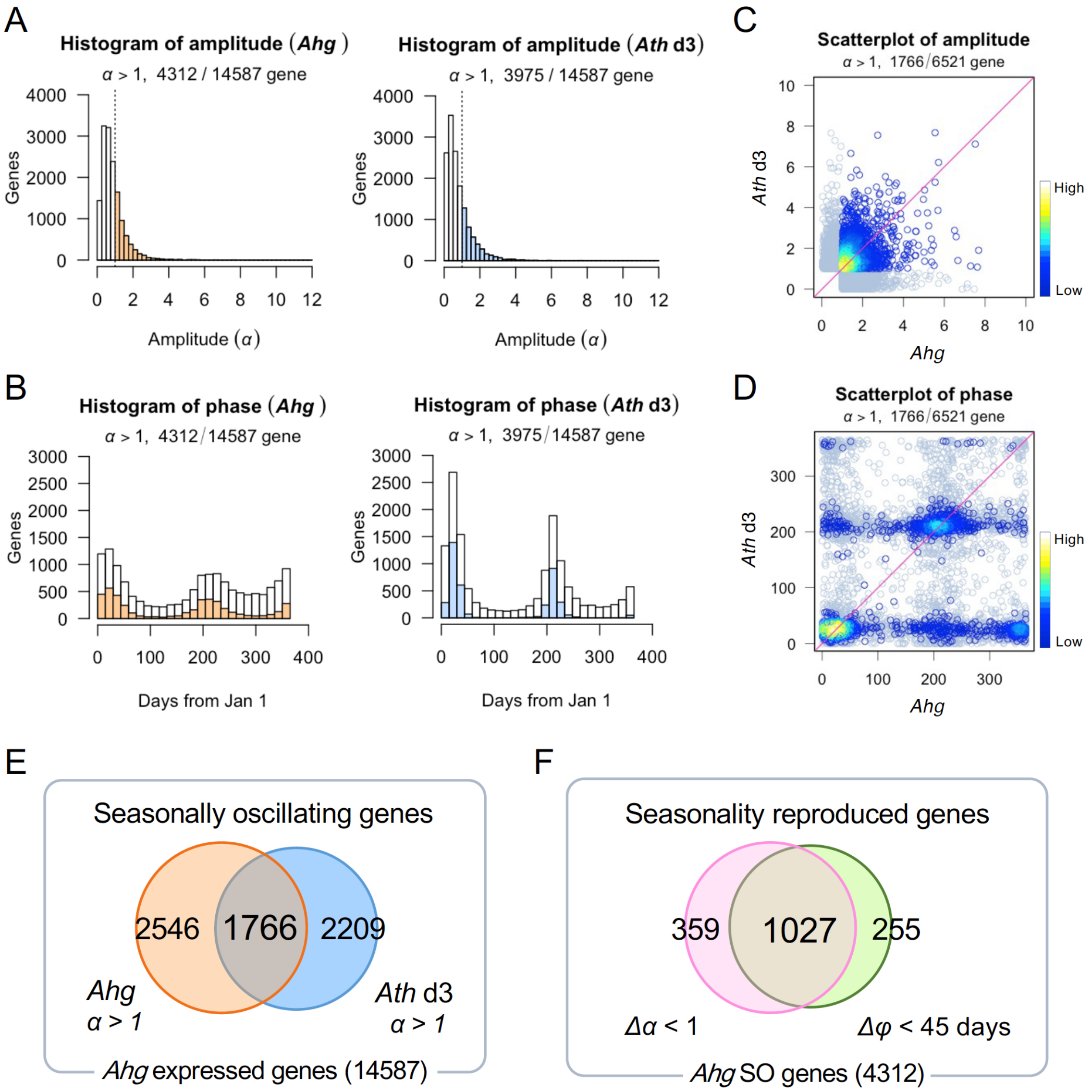

**Fig. S6** Comparison of seasonal trends of gene expression in *Arabidopsis halleri* ssp. *gemmifera* (*Ahg*) and *Arabidopsis thaliana* (*Ath*) grown in the SGCmini for 1 d. "$\alpha$" and "$\varphi$" indicate the amplitude and the phase of fitted cosine curves. (A) Histograms of the amplitude of expressed genes. Left: *Ahg*. Right: *Ath*. (B) Histograms of the phase of expressed genes. Left: *Ahg*. Right: *Ath*. (C) Scatter plot of the amplitude of *Ahg* (vertical axis) and *Ath* (horizontal axis). Only genes with amplitude > 1 in both species were colored by the local density of points. (D) Scatter plot of the phase of *Ahg* and *Ath*. Coloring conditions are the same as (C). (E) Venn diagram of *Ahg* and *Ath* seasonally oscillating (SO) genes. Left circle: *Ahg*. Right circle: *Ath*. (f) Venn diagram of genes that passed the amplitude and phase criteria. Left circle: genes that passed the amplitude criteria ($\Delta\alpha$ < 1). Right circle: genes that passed the phase criteria ($\Delta\varphi$ < 45 d).

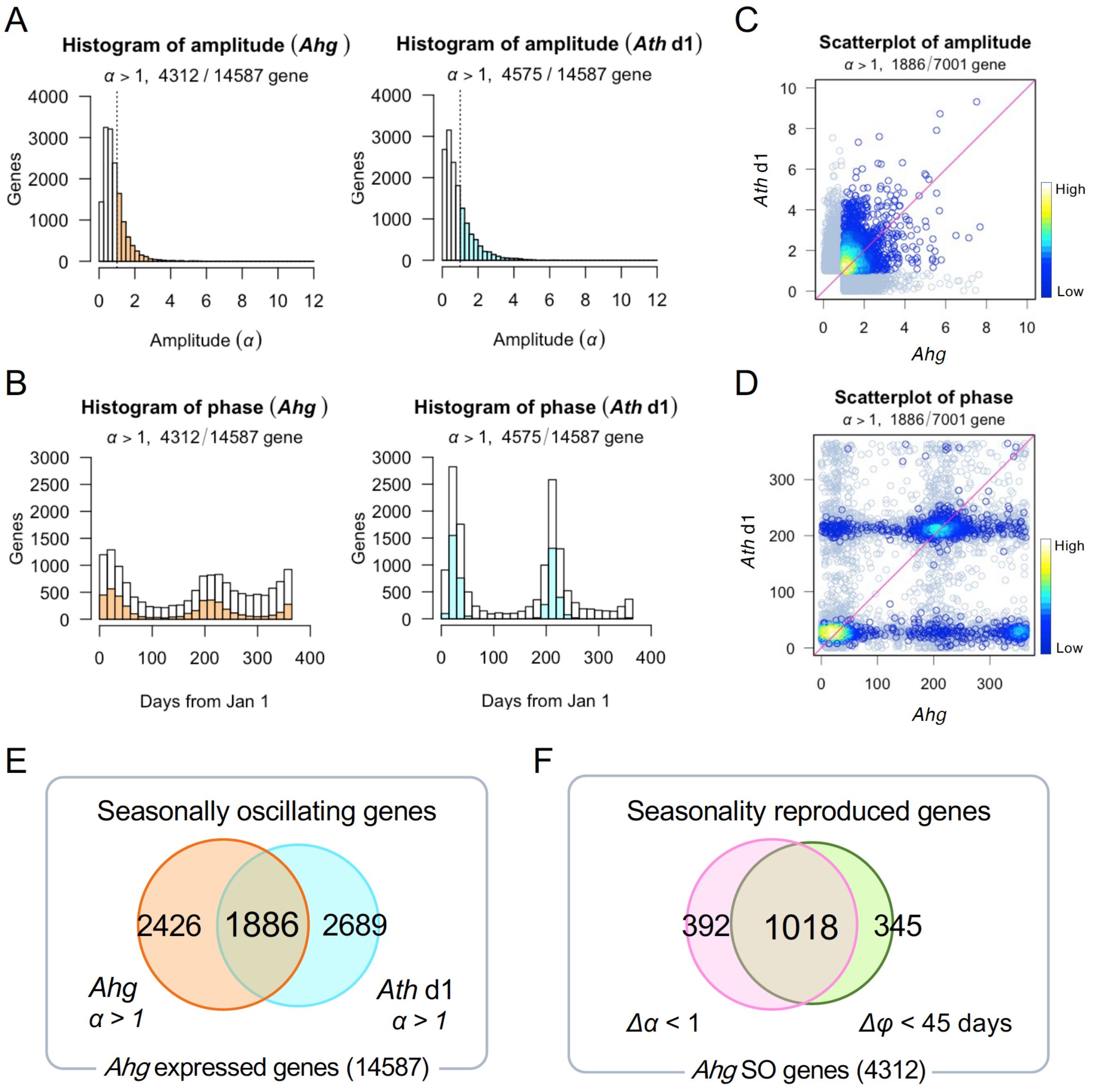

**Fig. S7** Anthocyanin regulatory and biosynthetic pathways and seasonal trend plots of *Arabidopsis halleri* ssp. *gemmifera* (*Ahg*) and *Arabidopsis thaliana* (*Ath*). (A) Fitted curve plots of early biosynthetic genes (EBGs) and late biosynthetic genes (LBGs). (B) Major transcriptional regulators that regulate EBGs. (C) Major transcriptional regulators that regulate LBGs. No ortholog of *MYB114* was found in *Ahg* by reciprocal BLAST. Observed value plots of each gene are shown in Fig. S8–10.

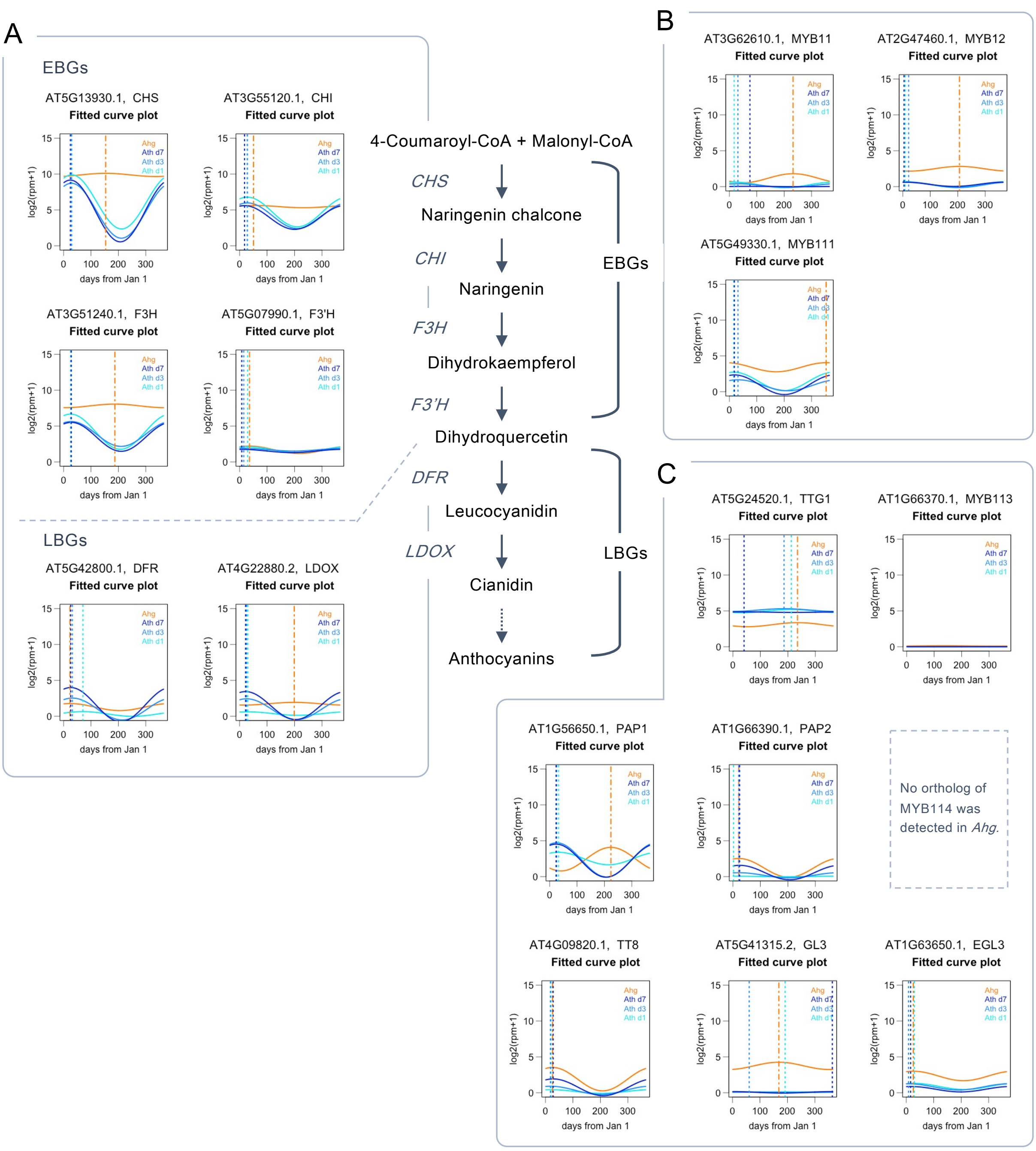

**Fig. S8** Seasonal expression plots of early biosynthetic genes (EBGs) and late biosynthetic genes (LBGs) in the anthocyanin biosynthetic pathway. From left, fitted cosine curve plot of *Arabidopsis halleri* ssp. *gemmifera* (*Ahg*) and *Arabidopsis thaliana* (*Ath*), observed value and fitted line plot of *Ahg* and *Ath* in the 7-, 3-, and 1-d cultures, respectively. (A) *CHS*. (B) *CHI*. (C) *F3H*. (D) *F3'H*. (E) *DFR*. (F) *LDOX*.

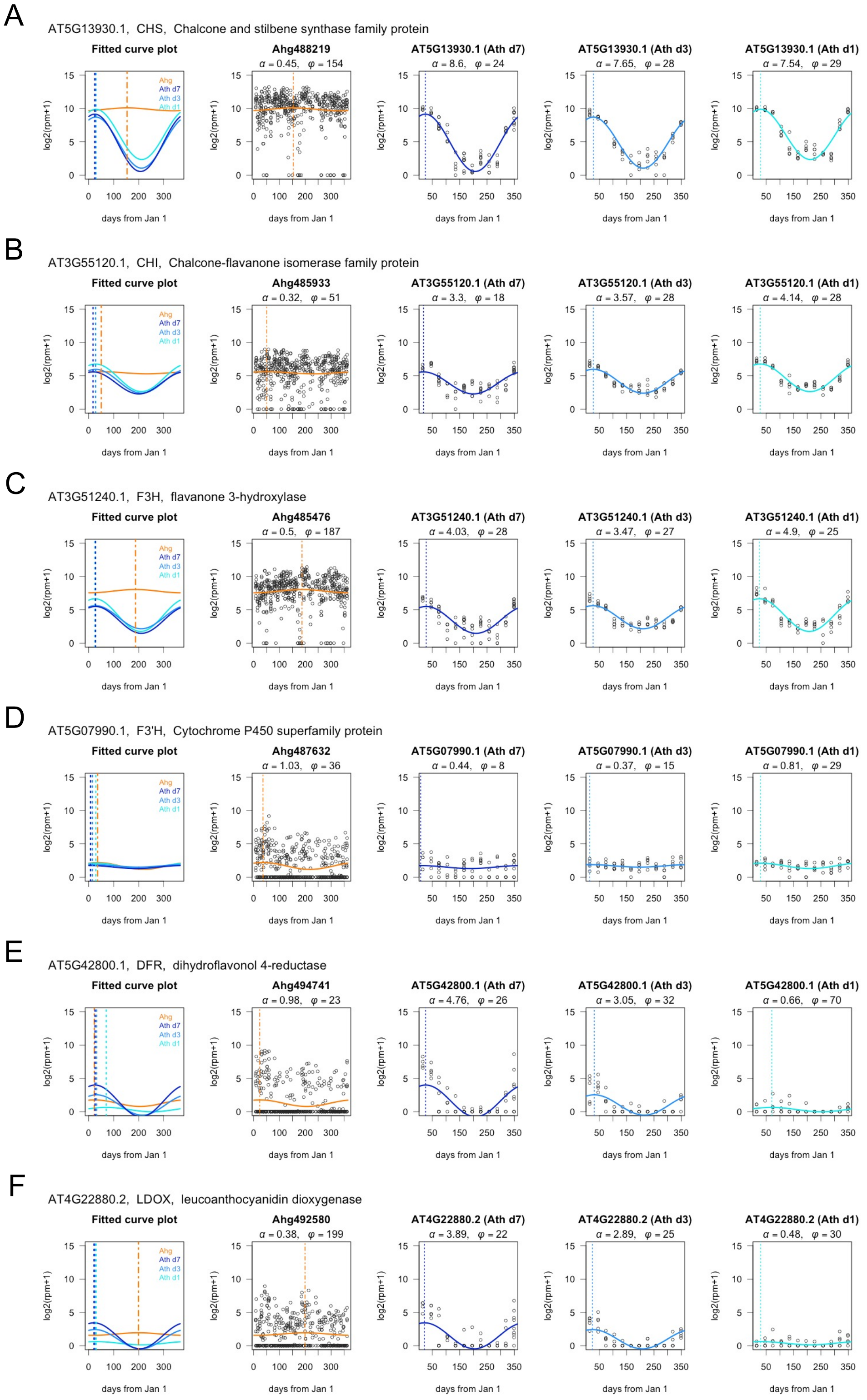

**Fig. S9** Seasonal expression plots of transcriptional regulators that regulate early biosynthetic genes (EBGs) in the anthocyanin biosynthetic pathway. From left, fitted cosine curve plot of *Arabidopsis halleri* ssp. *gemmifera* (*Ahg*) and *Arabidopsis thaliana* (*Ath*), observed value and fitted line plot of *Ahg* and *Ath* in the 7-, 3-, and 1-d cultures, respectively. (A) *MYB11*. (B) *MYB12*. (C) *MYB111*.

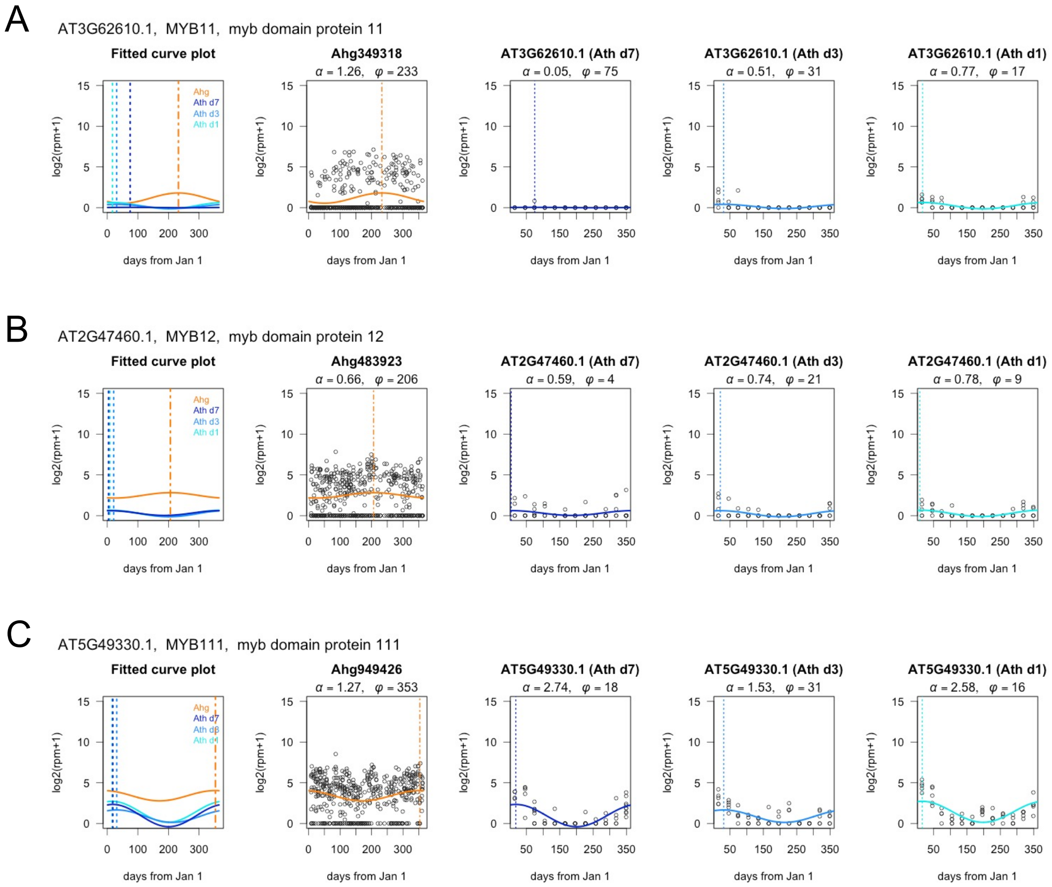

**Fig. S10** Seasonal expression plots of transcriptional regulators that regulate late biosynthetic genes (LBGs) in the anthocyanin biosynthetic pathway. From left, fitted cosine curve plot of *Arabidopsis halleri* ssp. *gemmifera* (*Ahg*) and *Arabidopsis thaliana* (*Ath*), observed value and fitted line plot of *Ahg* and *Ath* in the 7-, 3-, and 1-d cultures, respectively. (A) *TTG1*. (B) *MYB113*. (C) *PAP1*. (D) *PAP2*. (E) *GL3*. (F) *EGL3*.

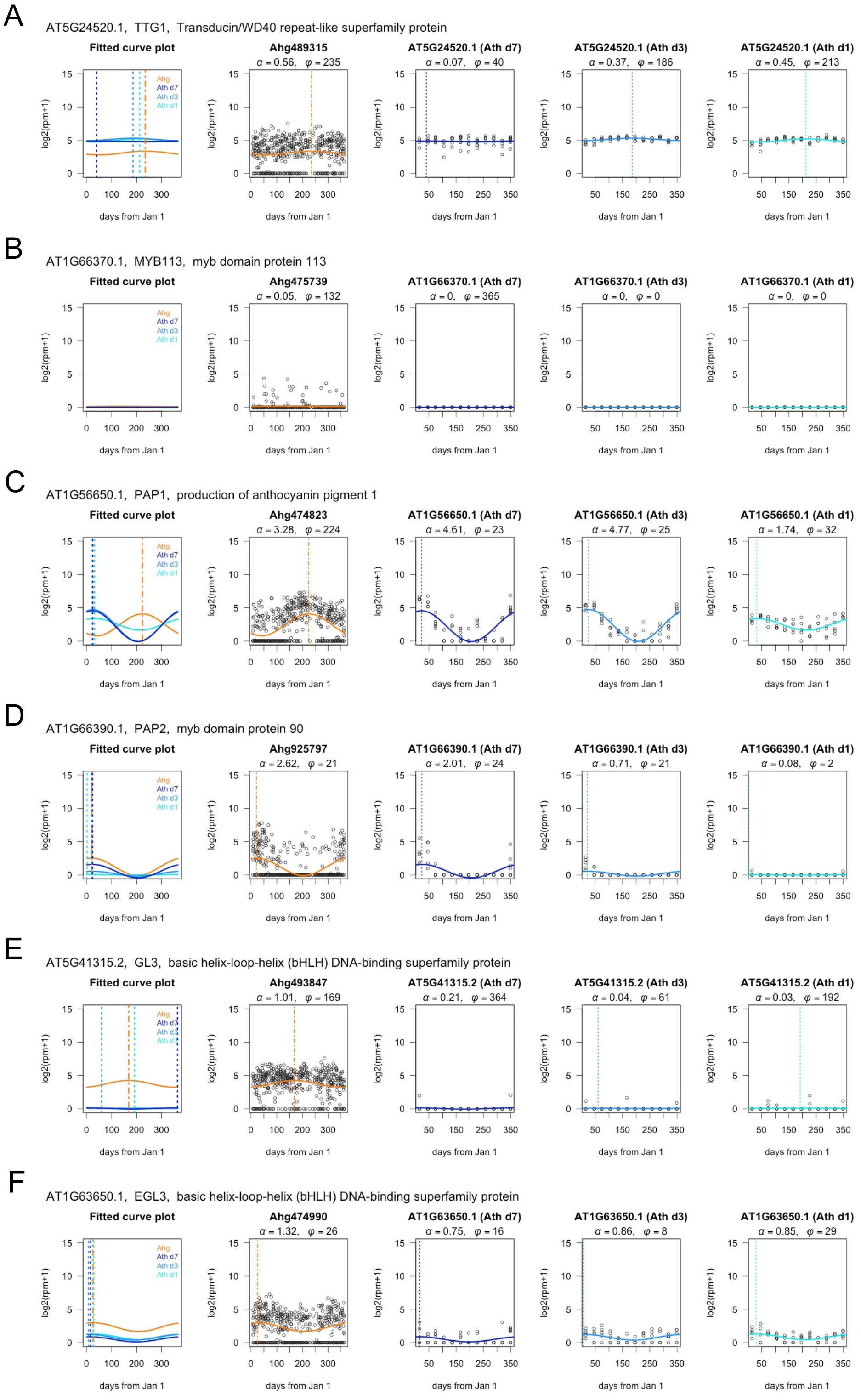

**Fig. S11** Comparison of gene expression patterns by length of pre-culture period. (A) Schematic diagram of culture conditions. To align the age of the plants with the 7-d culture condition, the pre-culture period was set to 3 d for the 1-d culture condition with short pre-culture. (B–D) Scatter plot of the difference in expression levels ($\log_2$(rpm+1)) of seasonally oscillating (SO) genes between August and January. (B) One-day culture (d1) versus 1-d culture with short preculture (s1). (C) One-day culture versus 7-d culture (d7). (D) One-day culture with short preculture versus 7-d culture. In all three cases, only SO genes were plotted. The local density of data points is represented using a color scale.

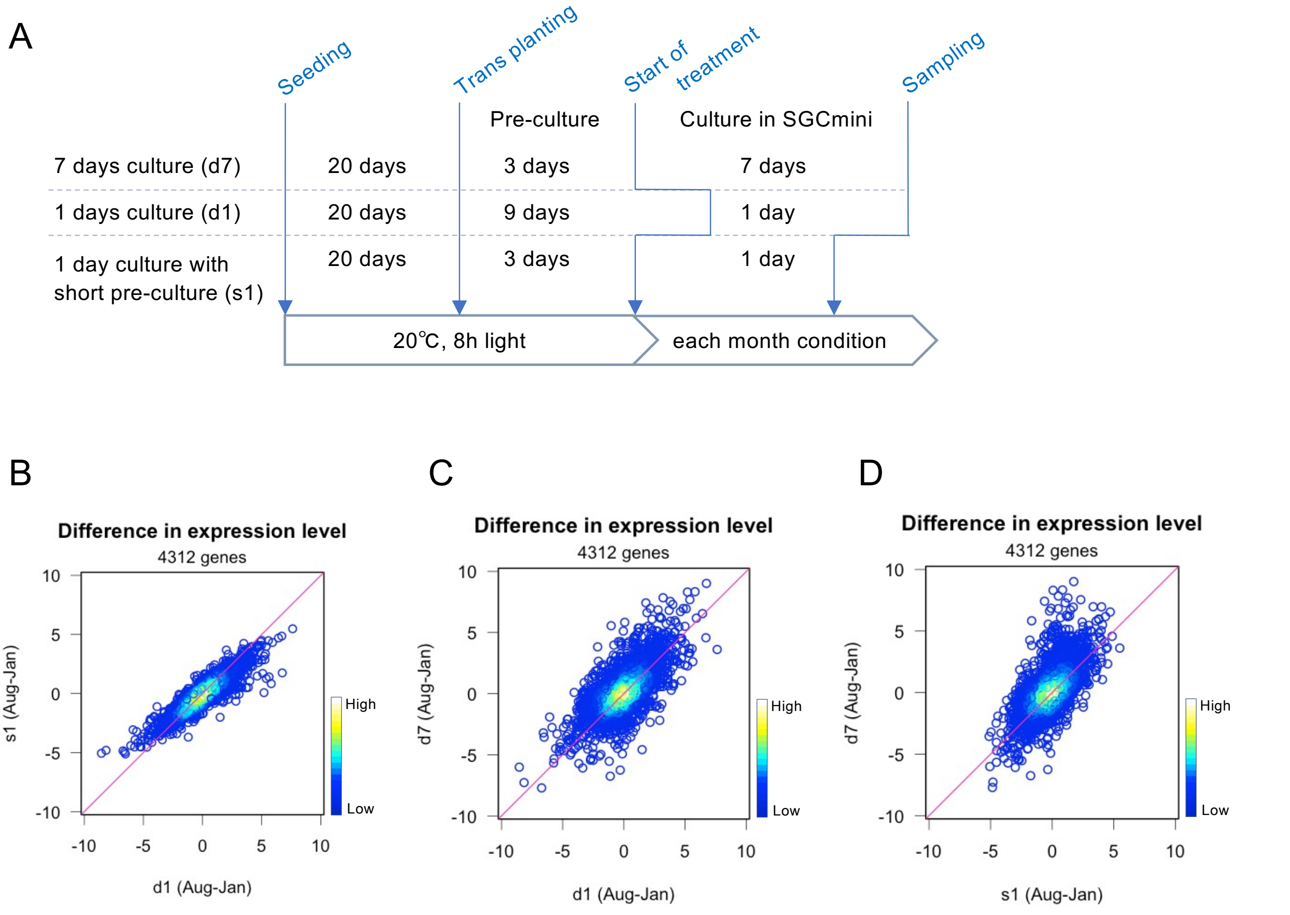

**Fig. S12** Enriched biological process gene ontology (GO) terms in SR genes for each condition. Circle size indicates the number of genes. Color scale indicates the adjusted *P* values in the GO enrichment analysis.

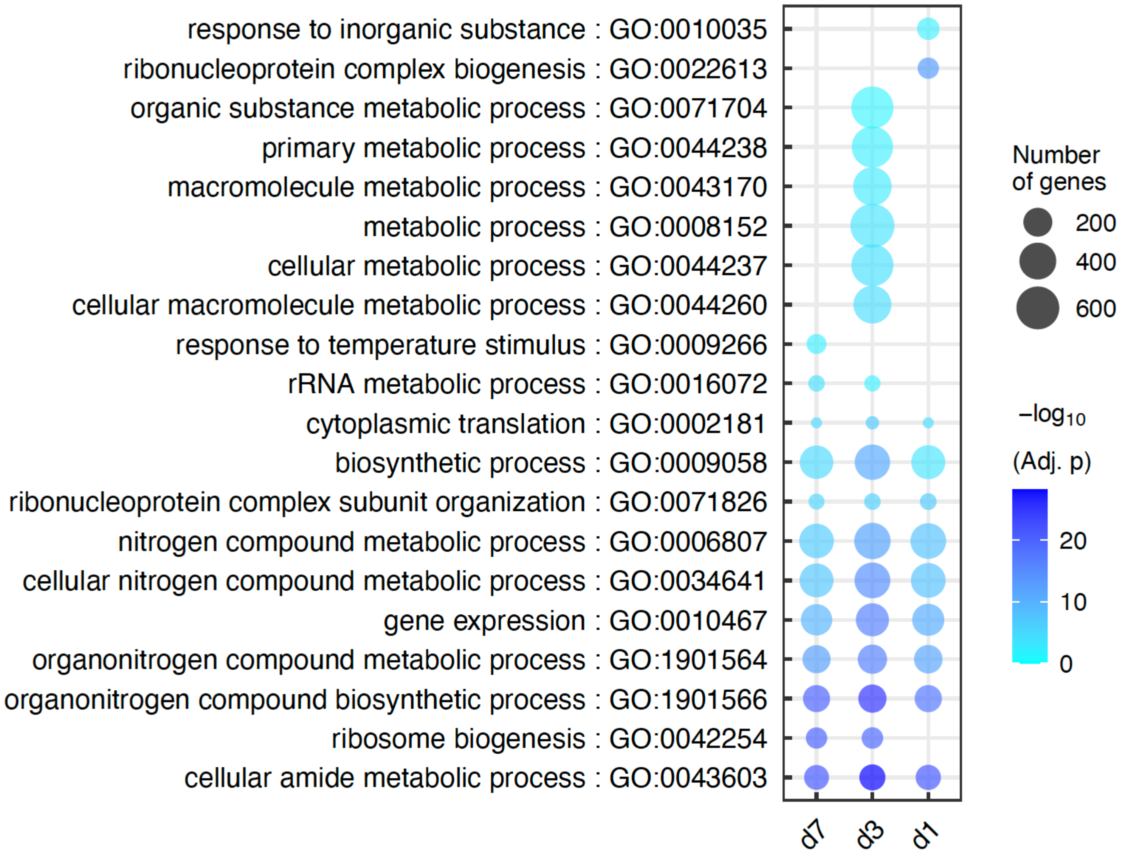
